# Mechanism of actin-dependent activation of nucleotidyl cyclase toxins from bacterial human pathogens

**DOI:** 10.1101/2021.06.27.450065

**Authors:** Alexander Belyy, Felipe Merino, Undine Mechold, Stefan Raunser

**Affiliations:** Department of Structural Biochemistry, Max Planck Institute of Molecular Physiology, Otto-Hahn-Str. 11, 44227 Dortmund, Germany; Department of Protein Evolution, Max Planck Institute for Developmental Biology, Max-Planck-Ring 5, 72076, Tübingen, Germany; Unité de Biochimie des Interactions Macromoléculaires, Département de Biologie Structurale et Chimie, Institut Pasteur, CNRS UMR 3528, Paris, France

## Abstract

Several bacterial human pathogens secrete nucleotidyl cyclase toxins, that are activated by interaction with actin of the eukaryotic host cells. However, the underlying molecular mechanism of this process which protects bacteria from self-intoxication is unclear. Here, we report structures of ExoY from *Pseudomonas aeruginosa* and *Vibrio vulnificus* in complex with their corresponding activators F-actin and profilin-G-actin. The structures reveal that in contrast to the apo state, two flexible regions become ordered and interact strongly with actin. The specific stabilization of these regions results in an allosteric stabilization of the distant nucleotide binding pocket and thereby to an activation of the enzyme. Differences in the sequence and conformation of the actin-binding regions are responsible for the selective binding to either F- or G-actin. This specificity can be biotechnologically modulated by exchanging these regions from one toxin to the other.

Other bacterial nucleotidyl cyclases, such as the anthrax edema factor and CyaA from *Bortedella pertussis*, that bind to calmodulin undergo a similar disordered-to-ordered transition during activation, suggesting that the allosteric activation-by-stabilization mechanism of ExoY is paradigmatic for all bacterial nucleotidyl cyclase toxins.

## Introduction

*Pseudomonas aeruginosa* and *Vibrio vulnificus* are human pathogens that cause nosocomial infections and seafood-related illnesses, respectively, with severity ranging from benign and local to systemic and life-threatening (1,2). A major virulence determinant of *P. aeruginosa* is the type 3 secretion system (T3SS), a cell wall associated nanomachine which transports a panel of effectors into eukaryotic target cells (3,4). One of such T3SS effectors is the nucleotidyl cyclase toxin exoenzyme Y (PaExoY) (5). It produces a supraphysiologic amount of 3’,5’-cyclic guanosine monophosphate (cGMP) and 3’,5’-cyclic adenosine monophosphate (cAMP) and thereby disorganizes cell signaling (6). At high concentrations, this results in the death of the cell when intoxicating cell cultures (7,8). At lower ExoY concentrations, which are more likely during chronical infection, the activity of PaExoY allows *P. aeruginosa* to hijack host immune response by a suppression of TAK1 and a decreased production of interleukin 1 (9,10). A recent report has established a link between the presence of PaExoY in clinical isolates and end-organ dysfunction in critically ill patients (11).

*V. vulnificus* also possesses an ExoY-like toxin (VvExoY), which acts as adenylate cyclase. It is essential for virulence in mice and is responsible for severe tissue damage (12). In contrast to PaExoY, VvExoY is not delivered by T3SS but as a module of MARTX (Multifunctional-Autoprocessing Repeats-in-ToXin). These large toxins combine toxic effector modules with their delivery apparatus in a single polypeptide chain (13).

Both PaExoY and VvExoY are catalytically inactive inside bacteria. However, once delivered into the target eukaryotic cell, the effectors bind to actin, resulting in more than 10,000-fold increase in nucleotidyl cyclase activity (14). A similar activation potency was observed in other members of nucleotidyl cyclase family, such as edema factor of *Bacillus anthracis* and adenylate cyclase domain of CyaA of *Bordetella pertussis*, which are activated by calmodulin. A series of structural studies revealed that interaction with calmodulin promotes a disordered-to-ordered transition in the catalytic center that is necessary for the efficient catalysis (15–17). However, it is not known if activation of nucleotidyl cyclases by actin follows the same molecular mechanism.

Here we report cryo-EM structures of PaExoY and VvExoY in complex with their activators F-actin and G-actin-profilin. Our structural data together with molecular dynamics simulations and enzymatic assays reveal that specific interactions between two independent regions of the toxins with F-, or G-actin lead to stabilization of the catalytic center of the toxins and high enzymatic activity. Thus, this mechanism appears to be general for the whole family of bacterial cyclase toxins.

## Results

### *P. aeruginosa* ExoY-F-actin complex

To decipher the activation mechanism of PaExoY, we solved the cryo-EM structure of the toxin with the substrate analog 3’-deoxyguanosine triphosphate (3’-dGTP) in complex with phalloidin-stabilized F-actin (Fig. 1A, Table S1). The map has an average resolution of 3.2 Å (Fig. S1, Table S1) and allowed us to build an almost full atomic model of the complex. PaExoY binds to F-actin in a 1:1 stoichiometry, which is consistent with our previous biochemical experiments (14). The absence of density between consecutive PaExoY monomers along the filament shows that there is no interaction between them.

**Fig 1.**
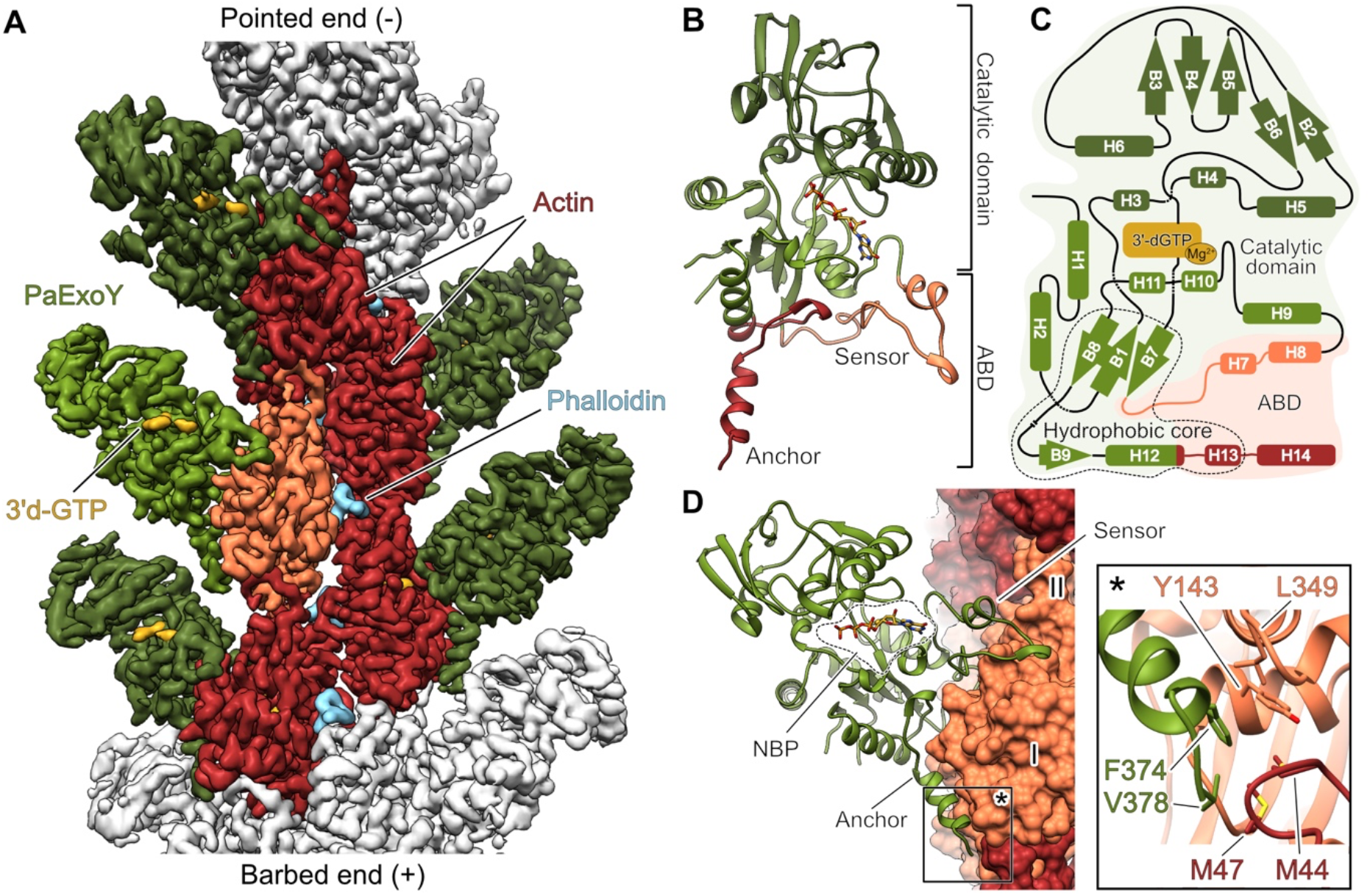
Cryo-EM structure of PaExoY in complex with F-actin. (A) The postprocessed map of the PaExoY-F-actin complex. The central actin subunit is colored in orange. Its surrounding 4 neighbors are shown in dark red. (B) Domain organization of PaExoY. (C) Schematic representation of the secondary structure elements of PaExoY. (D) Overview of the PaExoY-F-actin interface. Actin subdomains are shown in surface representation and are marked by roman numerals. ABD – actin-binding domain, NBP – nucleotide-binding pocket.

The overall structure of PaExoY bound to F-actin (Fig 1B and C) is similar to the structure of PaExoY in the apo state (18) (Fig S2A). It is composed of a smaller actin-binding domain (ABD) that is connected via a hydrophobic core (Fig S2B) to a larger catalytic domain with the nucleotide binding pocket (NBP) in its center. In contrast to the crystal structure of the apo state which is based on an enzyme that was treated by limited *in situ* proteolysis to remove flexible regions, the NBP and ABD are resolved in the PaExoY-F-actin cryo-EM map (Fig S2A). The ABD adapts to the surface of the filament and forms two extensive contacts with subdomains I and III of actin (Fig 1D, Fig S2A and C).

The central contact region, which we refer to as sensor (amino acids Ile-222 – Glu-258), interacts with the nucleotide-binding region and subdomain I of actin. At the end of this interface sits a loop with three aromatic residues that create a scaffold for the formation of a salt bridge between Asp-247 of PaExoY and Arg-95 of F-actin (Fig S2C). We performed a thorough mutational analysis of this region and either depleted the aromatic residues or the salt bridge and tested the binding affinity and enzymatic activity *in vitro* as well as toxicity in yeast (Fig S2D-G). Interestingly, these mutations did not affect the affinity of the toxin to F-actin but tremendously impaired its enzymatic activity and toxicity. Therefore, while this region of the sensor does not strongly contribute to the affinity of the complex, it plays an essential role in toxin activation.

The other PaExoY-F-actin interface is formed by the C-terminal region of PaExoY, that we call anchor (amino acids Lys-347 – Val-378). It is composed of a loop and a helix, interacting with a hydrophobic groove on the surface of subdomain I of actin (Fig 1D). Interestingly, Phe-374 and Val-378 at the tip of the helix extend over to the hydrophobic interface between two neighboring actin subunits (Fig 1D). Indeed, deletions of the 5 C-terminal residues in PaExoY completely abolished binding to F-actin and activity (Fig S2D-G), suggesting that unlike the sensor, the anchor largely determines the affinity of the toxin to Factin. Since this pocket exists only in filamentous actin, Phe-374 and Val-378 are likely key residues that provide specificity for F-actin.

### Mechanism of PaExoY activation

The NBP is conserved between members of the nucleotidyl cyclase toxins (5,18,19). Our structure of PaExoY in complex with F-actin reveals for the first time the atomic details of the active center of an actin-activated nucleotidyl cyclase. In the NBP, Arg-63, Lys-81 and Lys-88 interact with phosphate groups of 3’d-GTP and orient the nucleotide for hydrolysis (Fig 2A). Phe-83, Glu-258, Ser-292, Asn-297 and Pro-298 create a pocket that harbors the base of the nucleotide (Fig 2A). Finally, a third group of residues, including Asp-212, Asp-214 and His-291 are localized in the proximity of the ribose of 3’d-GTP. Since His-291 is localized ~ 4 Å away from the ribose, it probably acts as a nucleophilic base for deprotonation of 3’OH group of GTP (Fig 2A). There is an ongoing debate whether one or two Mg^2+^ ions are required for the efficient catalysis of bacterial nucleotidyl cyclases (15,20). In our structure, we see only a weak density at the position, where Mg^2+^ presumably coordinates the β and *γ* phosphates of the nucleotide, but we observe a clear density for a metal ion, likely Mg^2+^, that is coordinated by Asp-212, Asp-214 and His-291.

**Fig 2.**
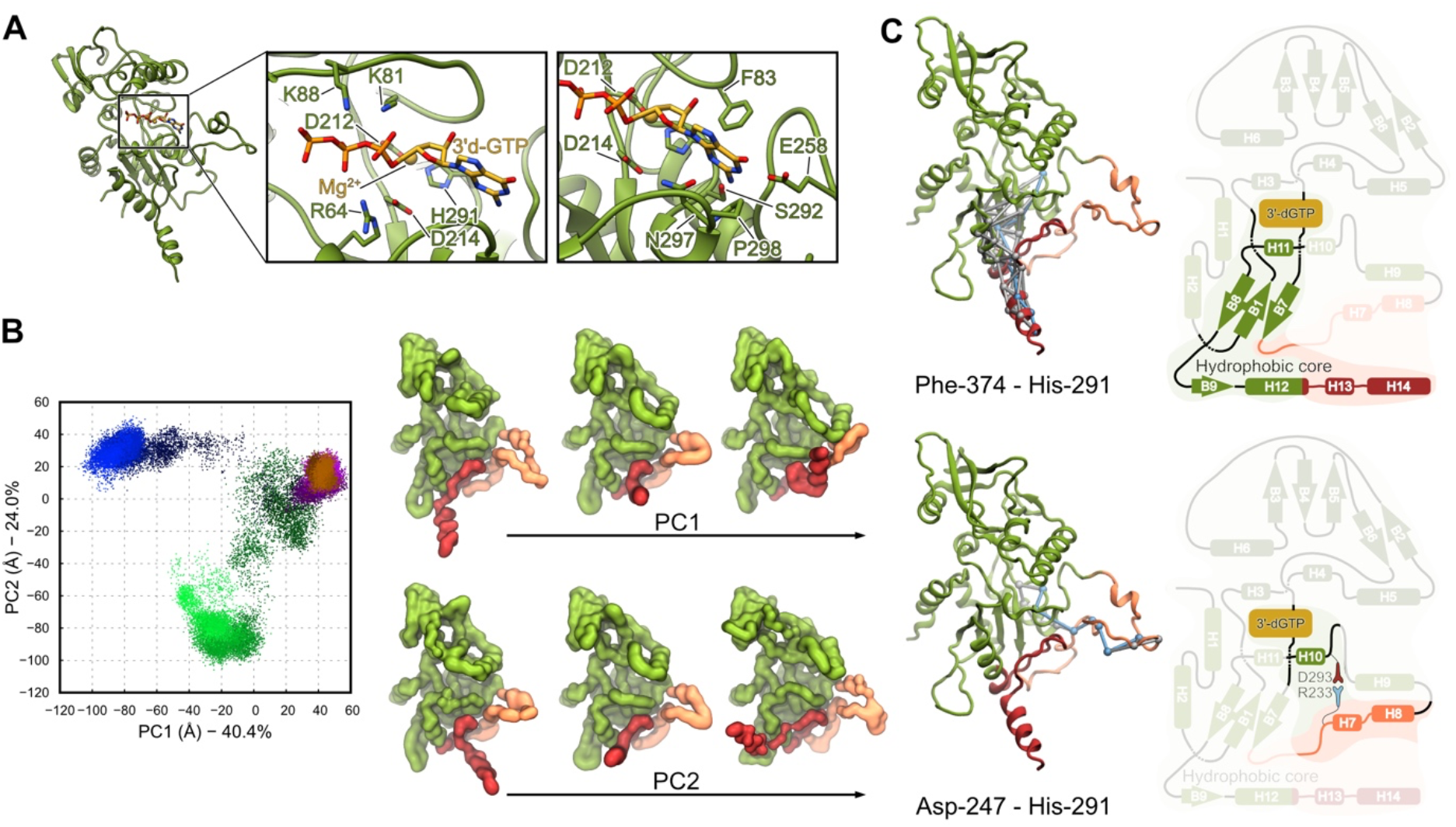
Effect of F-actin binding on the structure of PaExoY. (A) Atomic model of the PaExoY with focus on the nucleotide-binding pocket (NBP). (B) Principal component (PC) analysis of the different simulations. Each point represents a conformation projected onto the first two principal components. For each trajectory, time progresses as the color goes from dark to bright. The structures on the right from the plot show the nature of the variation along each component. (C) Collection of the paths (white) with the shortest possible path (light blue) connecting His-291 at the active site of PaExoY with different regions interacting with actin.

Due to flexibility, the lower part of the NBP has been absent in the crystal structure of PaExoY in the apo state (18) (Fig S2A). Stabilization of the pocket induced by the interaction of PaExoY with F-actin would therefore explain the strong activation effect of F-actin on the toxin.

To further explore the mechanism of activation of PaExoY, we performed molecular dynamics (MD) simulations of the toxin with and without F-actin, starting from its actin-bound conformation. In the absence of F-actin, PaExoY quickly drifts away from its starting conformation, with particularly strong fluctuations in the ABD and in parts of the NBP (Fig 2B, Fig S3A and B, Movie S1). In strong contrast, in simulations of the PaExoY-F-actin complex, the toxin reproducibly maintains its activated conformation, even when no substrate is present at its active site (Fig 2B, Fig S3A and B, Movie S1). Overall, the calculations suggest that a part of PaExoY undergoes a disordered-to-ordered transition upon an induced fit binding to Factin which results in the activation of the enzyme.

To decipher how the activation is communicated between the peripheral ABD and the central NBP, we applied dynamic network analysis to the MD simulation results (21). With this method, amino acids are represented as nodes in a network. Only amino acids with stable interactions during the simulation are connected, and their connection strength (the edge weight) is proportional to the correlation between the motion of the two amino acids in the simulations.

The dynamic network analysis revealed that structural changes in the ABD are transmitted to the center of PaExoY via two distinct pathways. While changes at the anchor and at the adjacent part of the sensor are transferred to the NBP via the hydrophobic core, changes at the peripheral part of the sensor are directly transmitted to the NBP to helix 10 (Fig 2C). Arg-233 and Asp-293 are central residues of the latter pathway, forming a salt bridge between the sensor and the NBP (Fig. S3C). Single amino acid substitutions of these residues fully abolished toxin activation but not F-actin binding (Fig S3D, F and G), corroborating their direct involvement in the communication of the activation signal from the periphery to the center of the toxin. The same is true for the hydrophobic core. A point mutation in its core abolished toxin activation (Fig S3D, F and G), indicating that proper packing of this area is key for allosteric signal transduction.

### *V. vulnificus* ExoY-G-actin-profilin complex

In contrast to PaExoY, which is activated by F-actin, its homologues from *Vibrio nigripulchritudo* (VnExoY) (22) and the human pathogen *Vibrio vulnificus* (VvExoY) (Fig S4) are activated by G-actin, indicating a high level of specificity of ExoYs for their activator. To decipher the molecular basis for this specificity, we set out to solve structures of VvExoY and VnExoY. We started with VnExoY in complex with 3’d-ATP and G-actin. However, although we tested different constructs, actin isoforms and cryo conditions, strong anisotropy prevented us from obtaining a high-resolution reconstruction (Fig S5). We therefore tried the equivalent VvExoY complex instead, but faced the same problem. Often, additional components of a protein complex change the characteristics of a particle strongly enough that they distribute differently in the ice. We therefore decided to add the actin-binding protein profilin to the complex, which does not interfere with the activation of VvExoY (Fig S4). Since most of G-actin in cells is captured by profilin, this represents well the situation *in vivo*. The addition of profilin improved anisotropy and allowed us to determine the structure of the VvExoY-G-actin-profilin complex at 3.9 Å resolution (Fig S6, Table S1), which we used to build a complete model of the complex (Fig 3A).

**Fig 3.**
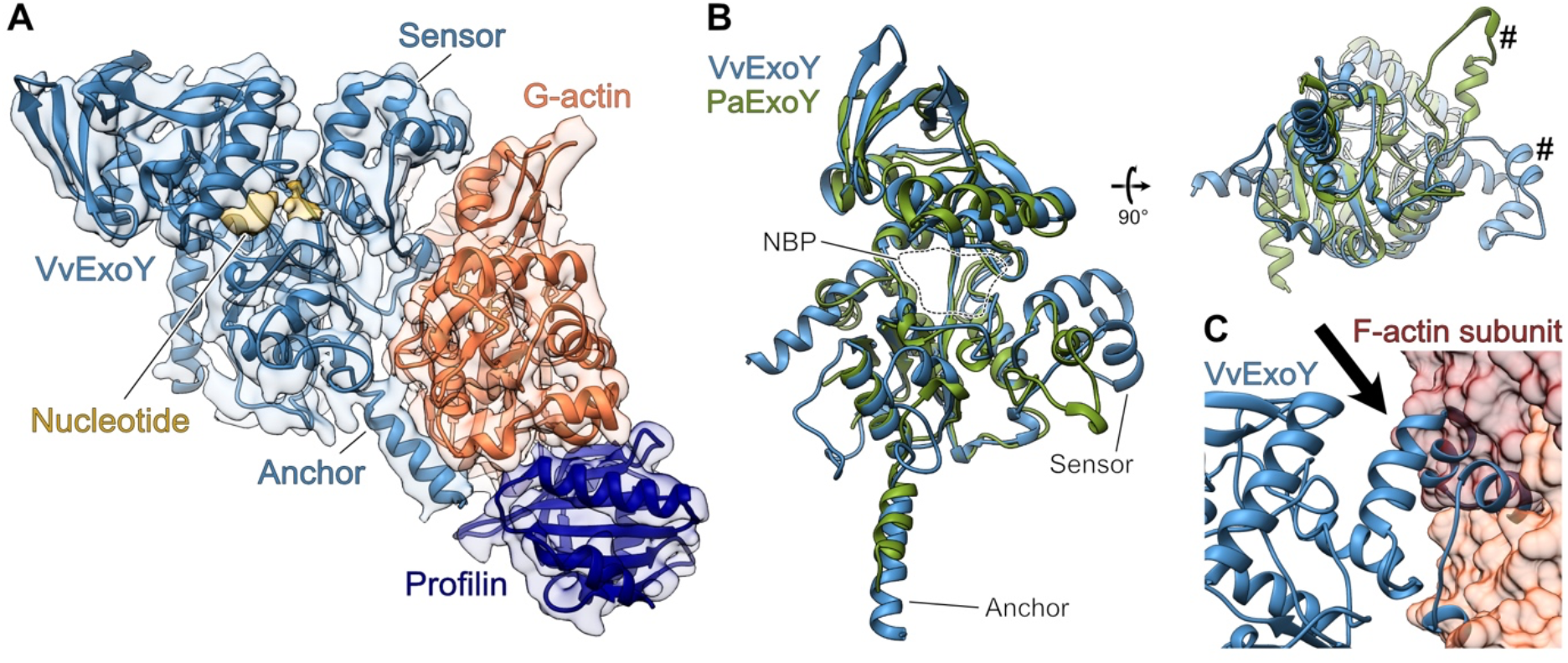
Cryo-EM structure of the VvExoY-G-actin-profilin complex. (A) An atomic model of the complex fit in the cryo-EM density that was combined from two maps (Fig S6C). (B) Alignment of atomic models of PaExoY and VvExoY. Hashtags indicate the major difference in the sensor of the toxins. NBP – nucleotide-binding pocket. (C) Docking of the VvExoY structure to F-actin shows a steric clash (arrow) between the central interaction region of VvExoY and the F-actin subunit.

The structure of VvExoY is very similar to PaExoY. However, there are three striking differences. One concerns the N-terminal helix which is rotated by 90° in the VvExoY structure. Since it is located at the opposite side of the VvExoY-G-actin interface, an influence of this region on the actin-binding properties are unlikely. The other two differences are found in the ABD. First, the C-terminal helix is extended by 6 residues in the case of VvExoY (Fig 3B). Interestingly, this does not increase the interface with G-actin in comparison to the PaExoY-F-actin complex since it protrudes away from actin.

The second difference in the ABD is more prominent, as the sensor of VvExoY is enlarged by 9 residues in comparison to PaExoY and rotated by ~90° allowing the ABD not only to establish contacts with subdomain I but also with subdomain II of G-actin. G-actin is in its expected globular conformation with a flexible D-loop. Since subdomain II is involved in intersubunit contacts in F-actin and parts of it are not accessible, the sensor of VvExoY would impair the binding to F-actin (Fig 3B and C), explaining the preference of this toxin for G-actin.

### Alteration of the G-/F-actin specificity of ExoY toxins

To corroborate that the ABDs are responsible for the selectivity of ExoY proteins for either F- or G-actin, we prepared PaExoY chimeras where we replaced parts the ABD with the corresponding regions in VnExoY (Fig 4, S7, Table S3). The construction of chimeras is in general a difficult task. Since these constructs often fail to express and function properly, we therefore prepared 15 different chimeras, of which 4 showed toxicity in yeast (Fig S8B). The change of the anchors resulted in a chimera (chimera A) that was equally activated by F- and G-actin (Fig 4). The affinity to F-actin was considerably reduced in comparison to the wildtype. Surprisingly, chimera B, where part of the sensor region that directly interacts with G-actin in VnExoY has been exchanged, still bound to F-actin and was preferably activated by the filament. This suggests that the altered sensor did not result in a steric clash with F-actin and corroborated that the anchor is very important for the selectivity of the toxin. When we exchanged the complete sensor plus a part of the catalytic center, the enzyme (chimera C) had a drastically reduced affinity to F-actin and strong preference for G-actin (Fig 3C). Interestingly, similar to VnExoY, chimera C produced only cAMP but not cGMP, indicating that the change in the NBP altered the substrate specificity of the enzyme (Fig 4C and D). When we exchanged both the anchor and sensor to the corresponding region of VnExoY, the resulting enzyme (chimera D) did not bind anymore to F-actin. Altogether, our structural and functional data on PaExoY and VvExoY complexes with actin and their chimeras show that the ABDs determine the selectivity of ExoY toxins for F- or G-actin. Thus, it should be possible to predict whether a certain ExoY is activated by F-actin or G-actin. To this end, we compared the sequences of all described ExoY-like proteins (14,18). After removal of repetitive and incomplete sequences, we ended up with 25 unique ExoY-like proteins (Fig S8). We analyzed the differences and similarities in the ABDs of these proteins and divided them into four groups. Based on the similarity to PaExoY or VvExoY, the first two groups represent likely F-actin and G-actin-activated ExoY-like proteins with the corresponding sensor and anchor regions. While 9 ExoYs recognize G-actin, only the ExoY from *Aeromonas salmonicida* and PaExoY seem to be activated by F-actin. Interestingly, the two other groups, encompassing 14 unique sequences, possess completely different activator-binding regions, suggesting an existence of a new class of bacterial cyclase toxins that are neither activated by calmodulin nor actin.

**Fig 4.**
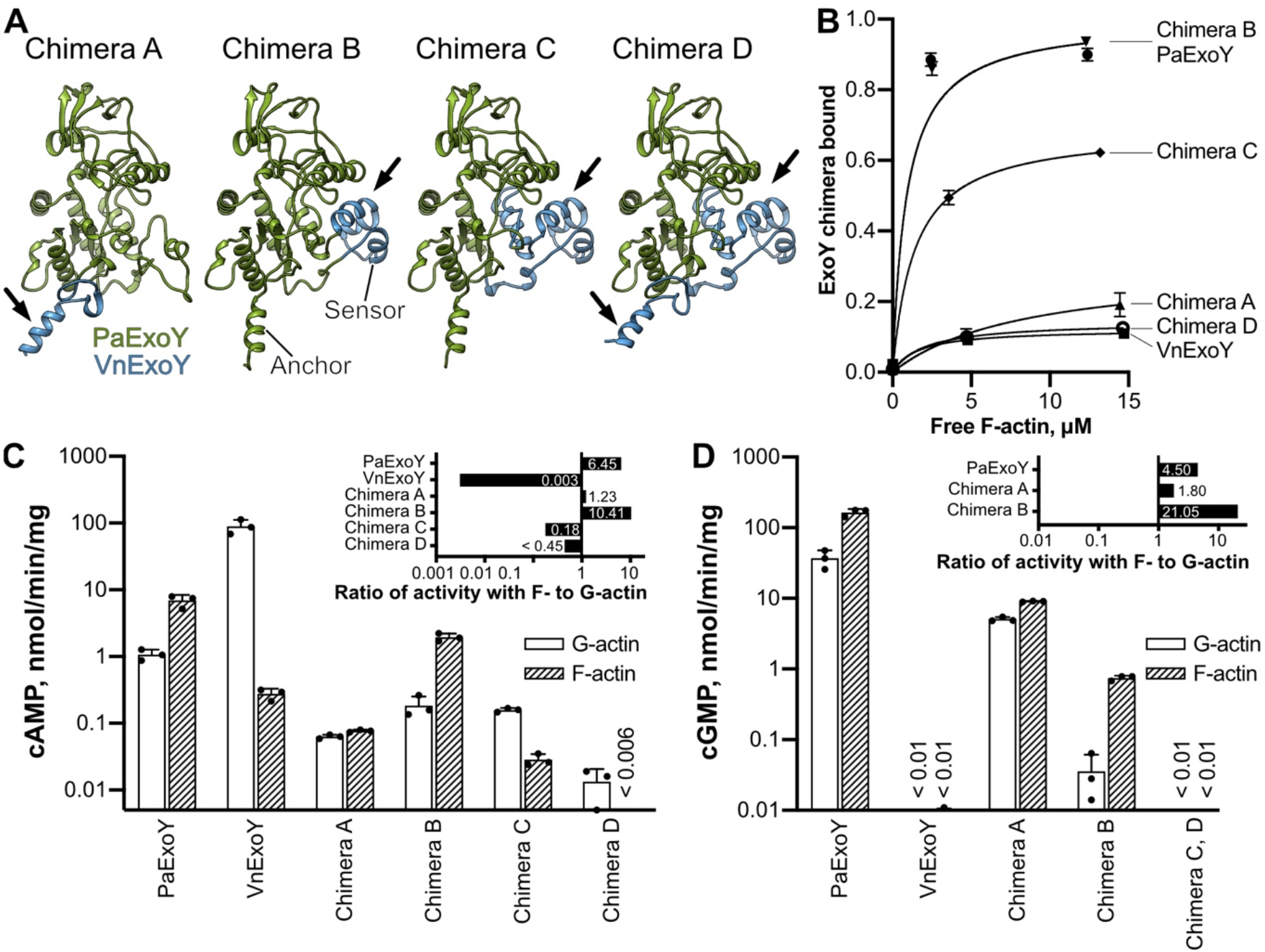
Chimera proteins of PaExoY and VnExoY. (A) Models of chimera proteins, where regions of PaExoY were swapped (arrows) with the corresponding parts of VnExoY obtained by homology modeling. (B) Cosedimentations of F-actin with 2.5 μM of PaExoY, VnExoY or the chimera proteins quantified by densitometry. (C) Adenylyl and (D) Guanylyl cyclase activity of nucleotidyl cyclases in the presence of 2 μM latrunculin-stabilized G- or phalloidin-stabilized F-actin, measured during 60 min of incubation. All other chimera proteins created in this study are presented in Fig S7 and Table S3. The error bars in panels B, C and D correspond to standard deviations of 3 independent experiments.

## Conclusion and Outlook

Our results allow us for the first time to describe the activation mechanism of actin-activated nucleotidyl cyclase toxins in molecular detail (Fig 5, Movie S2). In the absence of an activator, the toxins are inactive due to the flexibility of large parts of the enzymes. After translocation of the toxins into their eukaryotic target cells either by T3SS in the case of PaExoY or as part of MARTX toxin in the case of ExoYs from *Vibrio* species, they bind to F-actin or G-actin. During this process, the highly flexible ABD become ordered on the surface of actin, resulting in the stabilization of the whole toxin including the NBP. Thus, the enzyme is activated by an allosteric activation-by-stabilization mechanism. In the case of *P. aeruginosa*, the activity of the toxin results in sharp increase of levels of cGMP and cAMP that hijacks cell signaling in the immune cells allowing the pathogen to circumvent the host immune response (9,10). In the case of *V. vulnificus*, VvExoY produces cAMP which results in the disorganization of cell signaling and the promoting of inflammation in the infected area (12). Differences in the sequence and conformation of the ABD are responsible for the selective binding to either F- or G-actin. Notably, the specificity for the monomeric or filamentous actin can be biotechnologically modulated by exchanging the actin-binding regions from one toxin to the other. Based on our sequence analysis we identified a group of nucleotidyl cyclases that are likely neither activated by calmodulin nor actin. Future experiments will reveal the activator of these enzymes.

**Fig 5.**
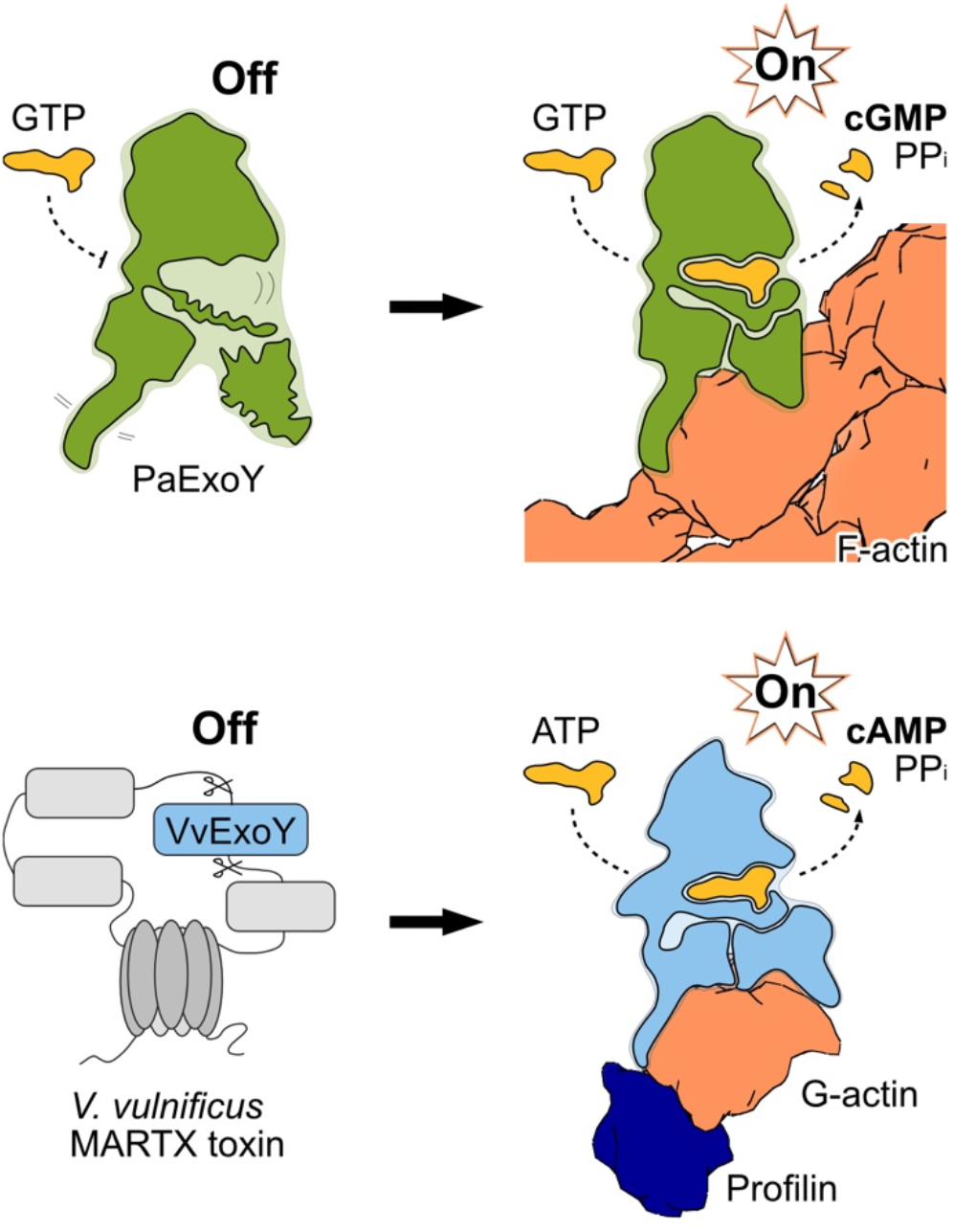
Activation of the actin-dependent nucleotidyl cyclase toxins from *P. aeruginosa* and *V. vulnificus*.

Other bacterial nucleotidyl cyclases, such as *B. anthracis* edema factor and CyaA from *B. pertussis*, that bind to calmodulin (Fig S3H), undergo a similar disordered-to-ordered transition during activation (15–17), suggesting that this mode of activation is paradigmatic for all bacterial nucleotidyl cyclase toxins. As many human pathogens contain nucleotidyl cyclases as effectors, we expect that our findings allow to better understand the molecular mechanism of pathogenesis of infection diseases and pave the road for the development of novel antidotes against poisons of microbial origin.

## Materials and methods

### Plasmids, bacteria and yeast strains, growth conditions

The list of oligonucleotides, constructions and strains used in this study can be found in the Supplementary data. *E. coli* strains were cultivated in LB medium supplemented with kanamycin or ampicillin. *S. cerevisiae* were grown on rich YPD medium or on synthetic defined medium (Yeast nitrogen base, Difco) containing galactose or glucose and supplemented if required with uracil, histidine, leucine, tryptophan, or adenine. *S. cerevisiae* strains were transformed using the lithium-acetate method (23). Yeast viability upon toxin expression under the galactose promoter was analyzed by a drop test (24). Analysis of protein expression in yeast was performed by total protein extraction (25), followed by SDS-PAGE, Western blotting, and incubation with anti-myc (9B11, CST) or anti-RPS9 serum (polyclonal rabbit antibodies were a generous gift of Prof. S. Rospert).

### Protein expression and purification

*V. vulnificus* ExoY is expressed as a module of MARTX toxins, which are post-translationally cleaved inside the host cell. As in the article we discuss exclusively the ExoY effector, we simplified amino acid numbering by designating amino acid 3229 of MARTX gene from *Vibrio vulnificus* strain BAA87 (GenBank: KJ131555.1) as the residue 1.

Fusion protein of *P. aeruginosa* ExoY, *V. nigripulchritudo* ExoY, *V. vulnificus* ExoY or their chimeras, and maltose-binding protein (MBP) were purified from *E. coli* BL21-CodonPlus(DE3)-RIPL cells possessing the plasmid listed in Table S2 according to the previously described protocol (26). *Vibrio nigripulchritudo* ExoY toxin without maltose binding protein was purified from *E. coli* BL21-CodonPlus(DE3)-RIPL cells possessing the plasmid pUM522 VnExoY according to the previously described protocol (22). Recombinant ExoY proteins were stored in the buffer containing 20 mM Tris pH 8 and 150 mM NaCl.

Rabbit skeletal muscle alpha-actin was purified as described previously (27) and stored in small aliquots at −80 °C. Human profilin-1 was a generous gift of Dr. J. Funk and Dr. P. Bieling that was purified according to the described protocol (28). Thymosin β4 was purchased from Sigma-Aldrich (reference SRP3324).

Mammalian cytoplasmic beta-actin was expressed in the C-terminal fusion with thymosin β4 and 10X-His-tag in insect cells BTI-Tnao38 in a similar strategy described by (29). As Cys-272 was shown to be prone to oxidation in aqueous solutions (30) we introduced a Cys272Ala mutation that is normally present in yeast actin gene. After expression, cells were resuspended in the lysis buffer containing 10 mM Tris pH 8, 50 mM KCl, 5 mM CaCl_2_, 1 mM ATP, 0.5 mM TCEP and cOmplete protease inhibitor (Sigma), and lysed using fluidizer. The supernatant after centrifugation was loaded on HisTrap FF crude (Thermo) equilibrated with the lysis buffer. After washing step with the same buffer, actin was eluted with a gradient of imidazole in lysis buffer. After overnight dialysis in G-buffer (5 mM Tris pH 8, 2 mM CaCl_2_, 0.5 mM ATP and 0.5 mM TCEP), actin was incubated with chymotrypsin for 20 min at 25°C to remove thymosin β4 and the following His-tag. After addition of PMSF to the final concentration of 0.2 mM to stop the cleavage reaction, the mixture was applied onto HisTrap FF column. The actin-containing flow-through was collected and polymerized overnight by addition of KCl and MgCl_2_ to the concentrations of 100 mM and 2 mM, respectively. The next day actin was span down at 210,000 g for 1 hour. The pellet was resuspended in G-buffer and dialyzed for at least 3 days against G-buffer. Finally, the protein was span down at 210,000 g for 1 hour, concentrated on 10 kDa cutoff Amicon columns, frozen in liquid nitrogen and stored at −80 °C.

### ExoY activity assays

Measurements of ExoY-dependent cAMP or cGMP synthesis *in vitro* were performed according to the previously described method (14) with modifications to avoid usage of radioactive materials. To start the reaction, 5 μl of 20 mM ATP or GTP solution was added to the 45 μl mixture of ExoY, 2 mM MgCl_2_ and G- or F-actin with phalloidin, latrunculin or actin-binding proteins, if indicated. After incubation at 30 °C, the reaction was stopped by the addition of 50 mm^3^ of Al_2_O_3_ powder that bind unreacted nucleoside triphosphates but does not interact with cyclic nucleotides. Following addition of 50 μl of TBS buffer, vortexing and centrifugation for 2 min at 15.000 g, light absorption of the supernatant was measured at the wavelength 252 or 259 nm for cGMP and cAMP, respectively. Amount of synthesized cNMP was then calculated using a calibration curve with defined concentration of cAMP or cGMP. In all reported experiments not more than 25% of the substrate was converted. Detection limit for each experiment was estimated as 0.1 optical density at the measured wavelength.

### Cosedimentation assays

Affinity of ExoY variants to F-actin was estimated using high-speed cosedimentation assays that were described previously (26).

### Cryo-EM sample preparation, data acquisition, processing and modelling of PaExoY-F-α-actin complex

Rabbit muscle F-actin was prepared as described previously (26). In brief, the freshly thawed protein was span down using TLA-55 rotor for 30 minutes at 150.000 g at 4 °C, and the G-actin-containing supernatant was collected. Then, the protein was polymerized by incubation in the buffer containing 120 mM KCl, 20 mM Tris pH 8, 2 mM MgCl_2_, 1 mM DTT and 1 mM ATP (F-buffer) in the presence of a twofold molar excess of phalloidin for 30 minutes at room temperature and further overnight at 4 °C. The next day, the actin filaments were pelleted using the same TLA-55 rotor for 30 minutes at 150.000 g at 4 °C and resuspended in the F-buffer. 15 minutes before plunging, F-actin was diluted to 2 μM, mixed with 4 μM of PaExoY, 2 mM 3’-deoxyguanosine-5’-triphosphate (3’-dGTP, Jena Bioscience) and 4 mM MgCl_2_. Shortly before the plunging, Tween-20 was added to the sample to a final concentration of 0.02% (w/v) to improve the ice quality. Plunging was performed using the Vitrobot Mark IV system (Thermo Fisher Scientific) at 13 °C and 100% humidity. 3 μl of sample were applied onto a freshly glow-discharged copper R2/1 300 mesh grid (Quantifoil), blotted for 8 s on both sides with blotting force −20 and plunge-frozen in liquid ethane.

The dataset was collected using a Krios Titan transmission electron microscope (Thermo Fisher Scientific) equipped with an XFEG at 300 kV using the automated data-collection software EPU (Thermo Fisher Scientific). Five images per hole with defocus range of −0.4 --3.5 μm were collected with the Falcon III detector (Thermo Fisher Scientific) operated in linear mode. Image stacks with 40 frames were collected with the total exposure time of 1.5 sec and total dose of 93 e^-^/Å^2^. 12437 images were acquired and 8663 of them were used for processing. Motion correction and CTF estimation have been performed in CTFFIND4 (31) and MotionCorr2 (32) during image acquisition using TranSPHIRE (33). The filament picking was performed using crYOLO (34). On the next step, 2.25 million helical segments were classified in 2D using ISAC (35) to remove erroneous picks. The remaining 1.85 million particles were used in the first 3D refinement in Meridien (36) with 25Å low-pass filtered Factin map as initial model and with a spherical mask with a diameter of 308 Å, followed by a local 3D refinement with a wide mask of the shape of the PaExoY-F-actin complex. On the next step, per-particle CTF-refinement was performed using 3D refinement parameters and the 3D reconstruction in Sphire, followed by removal of the segments with defocus value lower than −3 μm and recentering of the remaining 1.71 million particles according to the parameters calculated in the previous 3D refinement. Following the third round of 3D refinement in Sphire, Bayesian polishing and 3D classification were performed in Relion (37). The latter step was performed with a mask covering three actin and one PaExoY subunit in order to remove particles that do not possess bound PaExoY in the particular position. Indeed, 176067 segments with no PaExoY density were removed and the remaining 1.54 million were introduced into the final round of 3D refinement with a tight mask covering actin subunits and ExoY subunits. The final reconstruction map was postprocessed using 3D local filter based on the local resolution of the map in Sphire. Maps presented on Fig 1A and Movie S2 were postprocessed using DeepEMhancer (38). The overall processing, FSC curves and local resolution maps are available in the Fig S2.

To build a model of PaExoY-F-actin complex, we performed flexible fitting of the PaExoY partial crystal structure (5XNW, (18)) into the EM density using iMODFIT Chimera plugin (39). Then, we build the remaining 106 amino acids using Rosetta software (40,41) and added 3’-dGTP that was modelled with eLBOW (42). F-actin, ADP-P_i_ and Phalloidin were built as described previously for PDB 7AD9 (26). The model was further refined using ISOLDE (43) and Phenix (44). Schematic representation of the secondary structure elements of PaExoY for Fig 1C calculated in Chimera with H-bond energy cutoff −0.45 kcal/mol.

### Cryo-EM sample preparation, data acquisition, processing and model building of the VnExoY-G-α-actin complex

The freshly thawed rabbit muscle actin was span down using TLA-55 rotor for 30 minutes at 150,000 g at 4 °C, and the supernatant with G-actin was collected. Three μl of the sample containing 2.2 μM of VnExoY, 2 mM 3’-deoxyadenosine-5’-triphosphate (3’-dATP, Jena Bioscience), 4 mM MgCl_2_ and 2 μM G-actin, were applied onto a freshly glow-discharged copper R2/1 300 mesh grid (Quantifoil), blotted for 3 s on both sides with blotting force −3 and plunge-frozen in liquid ethane using the Vitrobot Mark IV system (Thermo Fisher Scientific) at 13 °C and 100% humidity.

The data collection was performed on a Krios Titan transmission electron microscope (Thermo Fisher Scientific) equipped with an XFEG at 300 kV and CS-corrector using the automated data-collection software EPU (Thermo Fisher Scientific). Four images per hole with defocus range of −1 --2.5 μm were collected with the Gatan K2 camera operated in counting mode. Image stacks with 64 frames were collected with total exposure time of 8 sec and total dose of 80 e^-^/Å^2^. 10,659 images were acquired and 9329 of them were used for processing. Motion correction and CTF estimation have been similarly performed in CTFFIND4 (31) and MotionCorr2 (32) during image acquisition using TranSPHIRE (33). Particle picking using crYOLO (34) resulted in 1.86 million particles that were used in 3D refinement in Meridien. For the latter, a 25Å low-pass filtered map of a single subunit of the PaExoY-F-actin complex was used as an initial model. After application of 2D shifts for recentering of the particles and removal of overrepresented views using Sphire, 2D classification was used to remove erroneous picks. The selected 1.1 million particles were subjected to three rounds of 3D refinement with particle restacking and particle polishing in Relion in between (37). After the second polishing step and another round of 2D classification, 3D refinement was performed with a tighter mask and initial model from the previous 3D refinement output. After the third round of particle polishing, the final round of 3D refinement was performed using Sidesplitter filter to reduce local overfitting due to preferred orientation of the particles in ice (45). The final reconstruction map was evaluated with 3D FSC tool (46) and postprocessed using DeepEMhancer (38). The processing overview with intermediate maps, angular distribution graphs and FSC curves is available in Fig S5.

### Cryo-EM sample preparation, data acquisition, processing and model building of the VvExoY-G-β-actin-profilin complex

The freshly thawed mammalian beta actin was span down using TLA-120 rotor for 20 minutes at 120,000 g at 4 °C, and the supernatant with G-actin was collected. A mixture containing G-actin at 18 μM, human profilin-1 at 24.5 μM, MBP-VvExoY at 18 μM, 3’-deoxyadenosine-5’-triphosphate (3’-dATP, Jena Bioscience) at 2 mM and MgCl_2_ at 4 mM was incubated for 15 minutes at room temperature. Shortly before plunging, the sample was diluted 5 times with 10 mM Tris pH 8 supplemented with 0.002% Tween-20. Then, the mixture was applied onto a freshly glow-discharged copper R2/1 300 mesh grid (Quantifoil), blotted for 3 s on both sides with blotting force −5 and plunge-frozen in liquid ethane using the Vitrobot Mark IV system (Thermo Fisher Scientific) at 13 °C and 100% humidity.

The data collection was performed on a Krios Titan transmission electron microscope (Thermo Fisher Scientific) equipped with an XFEG at 300 kV and CS-corrector using the automated data-collection software EPU (Thermo Fisher Scientific). Three images per hole with defocus range of −1.2 --2.5 μm were collected with the Gatan K3 camera operated in superresolution mode. Image stacks with 60 frames were collected with total exposure time of 2 sec and total dose of 60 e^-^/Å^2^. Following acquirement of 3773 movies without stage tilt, 3106 micrographs were imaged in an equal number using 10-, 20- or 30-degrees stage tilt. After visual inspection, 378 non-tilted and 183 tilted micrographs were removed. For the remaining images, motion correction and CTF estimation have been performed in CTFFIND4 (31) and MotionCorr2 (32) during image acquisition using TranSPHIRE (33). Particle picking using crYOLO (34) resulted in 706 thousand non-tilted and 543 thousand tilted particles that were used in independent 3D refinements in Meridien. For the latter, a 25Å low-pass filtered map of the simulated VnExoY-G-actin complex with docked profilin (PDB 2PAV) was used as an initial model. After application of 2D shifts for recentering of the particles, 2D classification was used to remove erroneous picks. The selected 222 and 130 thousand particles, non-tilted and tilted, respectively, were subjected to the second round of 3D refinement. Using 2D shift parameters obtained in the 3D refinement step, the particles were recentered and merged into one stack. After two more rounds of 3D refinement and particle polishing in Relion in between (37), the resulting 3.9 Å map was obtained and used for modeling of the VvExoY-G-actin complex. To compare the quality of the map with the density of the VnExoY-G-actin complex, the map was additionally evaluated with 3D FSC tool (46).

To further boost the profilin density, we performed additional alignment-free 3D classification in Relion, followed by 3D refinement in Meridien. The resulting 4.2 Å map was used to model the actin-profilin interface. The processing overview with intermediate maps, angular distribution graphs and FSC curves is available in Fig S6.

To build the model of VvExoY-G-actin-profilin complex, we first modelled VvExoY using tFold server (https://drug.ai.tencent.com/) and fit it into the EM density using iMODFIT Chimera plugin (39). Amino acids 1-38, 236-273, 399-432 that did not fit the experimental density were removed and remodeled using Rosetta software (40,41). The VvExoY model was then merged with profilin-G-actin complex from PDB 6NBW (47) and further refined using ISOLDE (43) and Phenix (44).

### Homology modeling

Homology models of the chimeric ExoY proteins were built using MODELLER (48). For this we used PaExoY and VvExoY as templates, but deleting the areas of the protein that should not be taken into account (e.g. PaExoY’s C-terminus in Chimera A).

### Molecular dynamics simulations

We used the cryo-EM model of the PaExoY-F-actin complex to set up classical molecular dynamics simulations of PaExoY in the presence or absence of F-actin. For free PaExoY, we took the coordinates of a single chain from the decorated filaments and solvated it with TIP3P waters 15 Å away from the furthest protein atom, using a truncated octahedron box. For the PaExoY-F-actin complex, we built a system composed of a single PaExoY molecule bound at the center of a filament made of 9 actin subunits. We then placed the complex into a TIP3P water box of dimensions 150×150×315 Å. KCl was then used to neutralize the systems and bring the ionic strength to 150 mM. All N-termini were acetylated. All simulations were prepared in CHARMM-GUI (49) using the CHARMM 36m forcefield (50). Simulations were run in NAMD 2.14 (51), using a time step of 2 fs. All bonds involving hydrogens were constrained to their equilibrium length. Short-range non-bonded interactions were truncated at 12 Å, with switching starting at 10 Å. Long-range electrostatics were treated using particle mesh Ewald (52). For each system we run 2 independent simulations starting from random velocities. The systems were equilibrated using the stepwise protocol summarized in Table S4, and then run for 400 ns.

Simulation results were analyzed using VMD (53) and Carma (54). The network analysis was performed using the NetworkView plugin of VMD (21).

## Acknowledgements

We thank O. Hofnagel and D. Prumbaum for assistance with data collection and maintaining the EM facility, S. Bergbrede for the excellent technical assistance, W. Linke and A. Unger (Ruhr-Universität Bochum, Germany) for providing us with muscle acetone powder, S. Rospert (University of Freiburg, Germany) for providing us anti-RPS9 serum, J. Funk and P. Bieling for providing us with profilin-1, and S. Pospich, Y. Belyi and D. Ladant for fruitful discussions. This work has been funded by the Max Planck Society (S.R.). A.B. was supported by a fellowship of the Humboldt foundation and an EMBO long-term fellowship.

## Author contribution

A.B. and S.R. designed the project. A.B. prepared cryo-EM specimens, collected and analyzed EM data, performed affinity and activity assays. A.B. built the atomic models with support from F.M. F.M. performed and analyzed MD simulations. A.B., and F.M. prepared figures. S.R. supervised the project. A.B., F.M. and S.R. wrote the manuscript with input from U.M.

## Data Availability

The coordinates for the cryo-EM structures of PaExoY-F-actin, VvExoY-G-actin-profilin and VnExoY-G-actin have been deposited in the Electron Microscopy Data Bank under accession number XXX. The corresponding molecular models for the PaExoY-F-actin, VvExoY-G-actin-profilin complexes have been deposited at the wwPDB with accession codes XXX.

