## Supplementary Figures for "Mechanism of actin-dependent activation of nucleotidyl cyclase toxins from bacterial human pathogens"

### Supplementary data

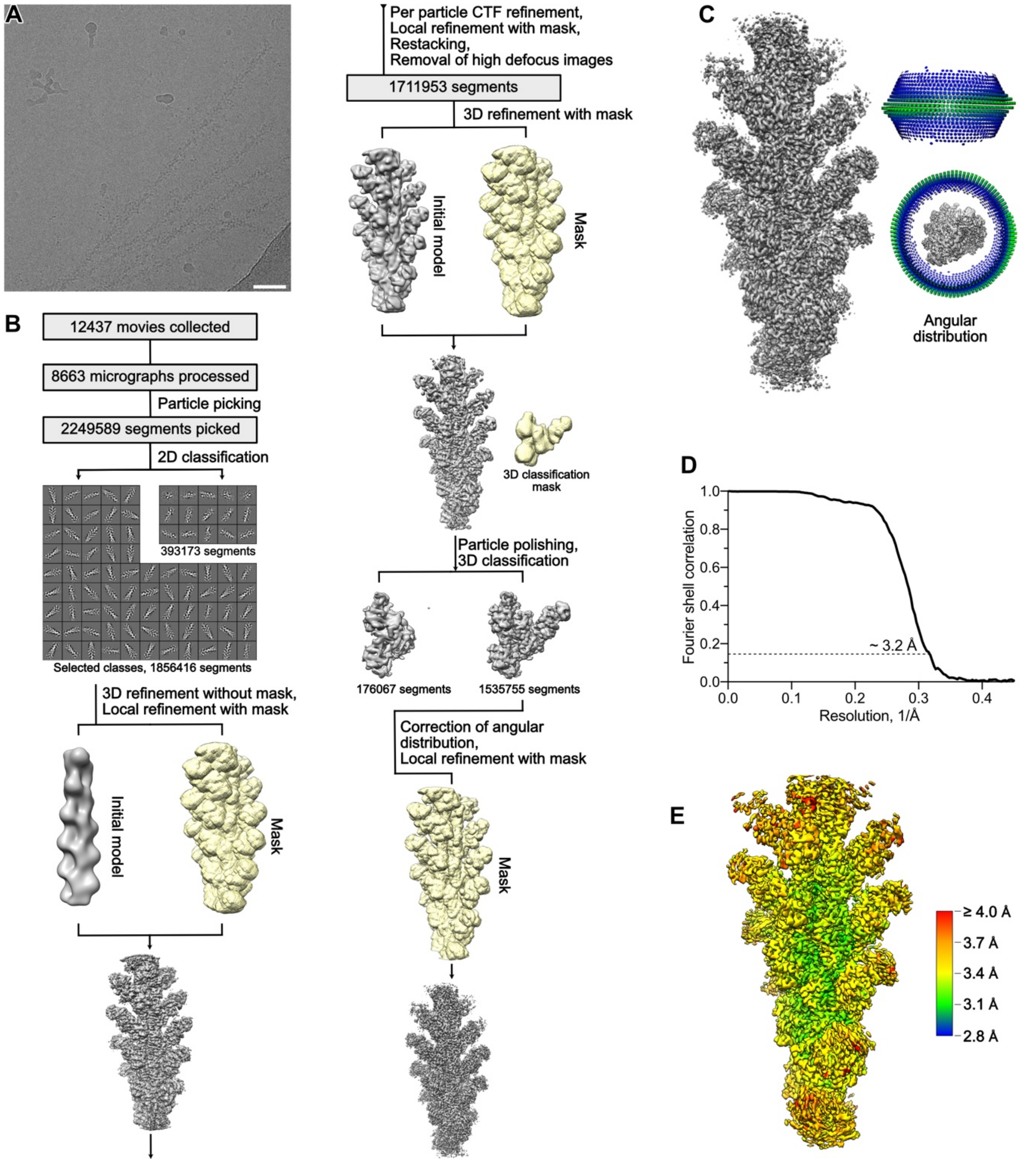

**Fig S1. Processing of the PaExoY-F-actin complex.** (A) An example cryo-EM micrograph. Scale bar 50 nm. (B) Processing overview. (C) The final postprocessed map filtered according to the local resolution, and its angular distribution. (D) Fourier shell correlation curve. (E) Local resolution gradient of the reconstruction.

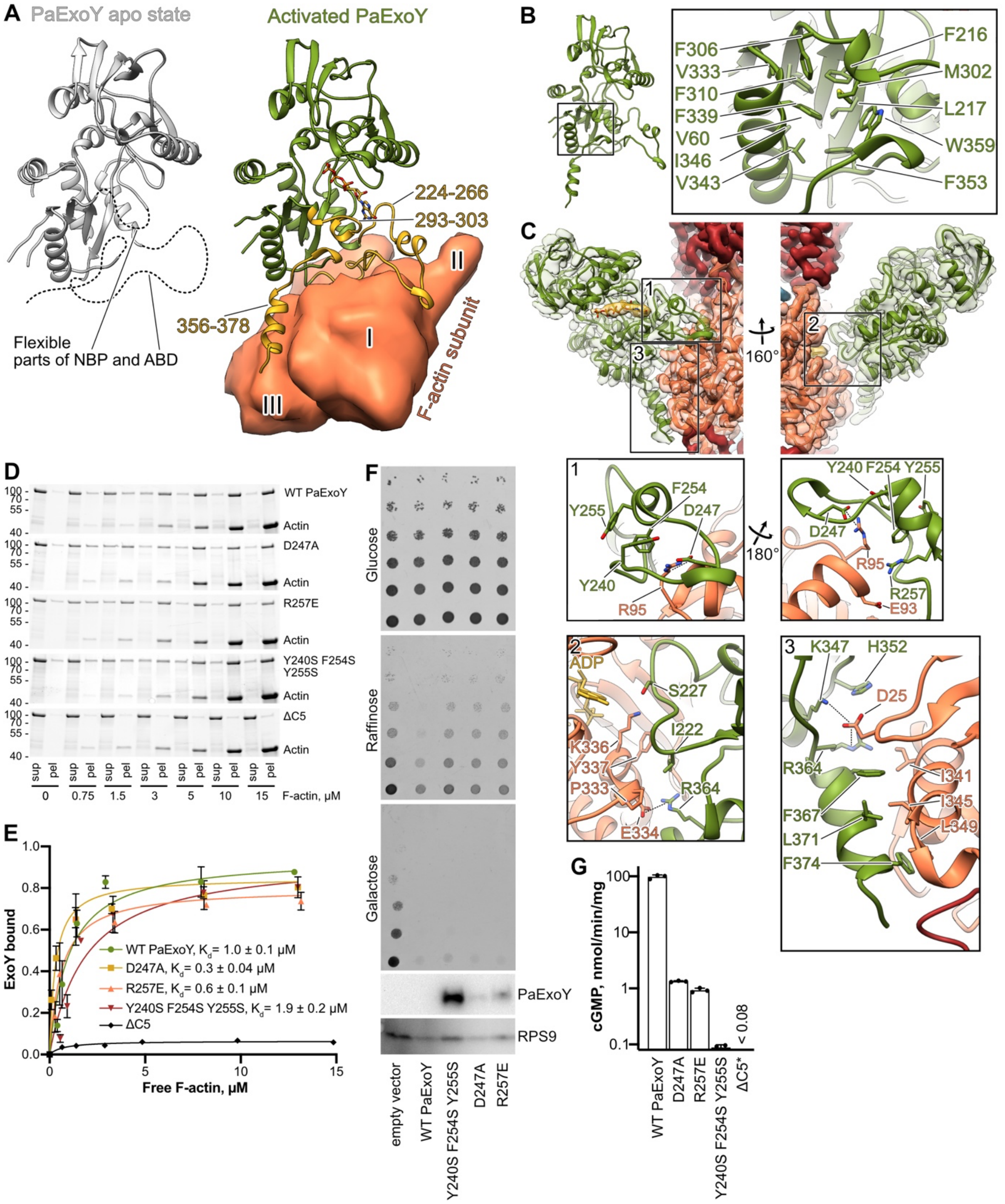

**Fig S2. PaExoY-F-actin complex.** (A) Comparison of the PaExoY apo state structure (1) to its activated form. Dashed lines indicate the missing regions in the apo sate. These regions are fully resolved in the PaExoY-F-actin structure and are colored in yellow. Actin subdomains are marked by Roman numerals. (B) Hydrophobic core of PaExoY (C) Overview of the interaction sites. Close-up views: 1 – distal part of the sensor with the hydrophobic scaffold of Tyr-240, Phe-254 and Tyr-255 that position Asp-247 and Arg-257 towards Glu-93 and Arg-95 of actin; 2 – Ile-222 at the proximal part of the sensor creates contacts with Tyr-337 and Pro-333 of subdomain III of actin; 3 – the anchor starts with a loop where positively charged Lys-347 and Arg-364 lock Asp-25 of actin, and continues in a long helix that creates extensive hydrophobic contacts with the subdomain I of actin. (D) Cosedimentation of F-actin and 2.5 μM PaExoY analyzed by SDS-PAGE. The upper band corresponds to PaExoY variants, and the lower band corresponds to actin. Representative stain-free gels are shown. (E) The fractions of PaExoY that cosedimented with F-actin were quantified by densitometry and plotted against F-actin concentrations. (F) Growth phenotype assay with *S. cerevisiae* expressing PaExoY variants under a strong galactose promoter in the experimental conditions with background (Glucose), low (Raffinose) or high (Galactose) toxin expression. Analysis of protein expression was performed by Western blot of cells grown on galactose-containing media with anti-myc (PaExoY) and anti-ribosomal protein S9 (RPS9) antibodies. (G) Activity of 30 ng of WT or 1 μg of PaExoY variants in the presence of 3 μM non-stabilized actin measured during 10 min of incubation. The error bars in panels E and G correspond to standard deviations of 3 independent experiments. ΔC5 is an PaExoY mutant with a deletion of 5 C-terminal amino acids. * - The measurement was performed previously (2). ABD – actin-binding domain, NBP – nucleotide-binding pocket. The uncropped gels and Western blots can be found in Fig S9.

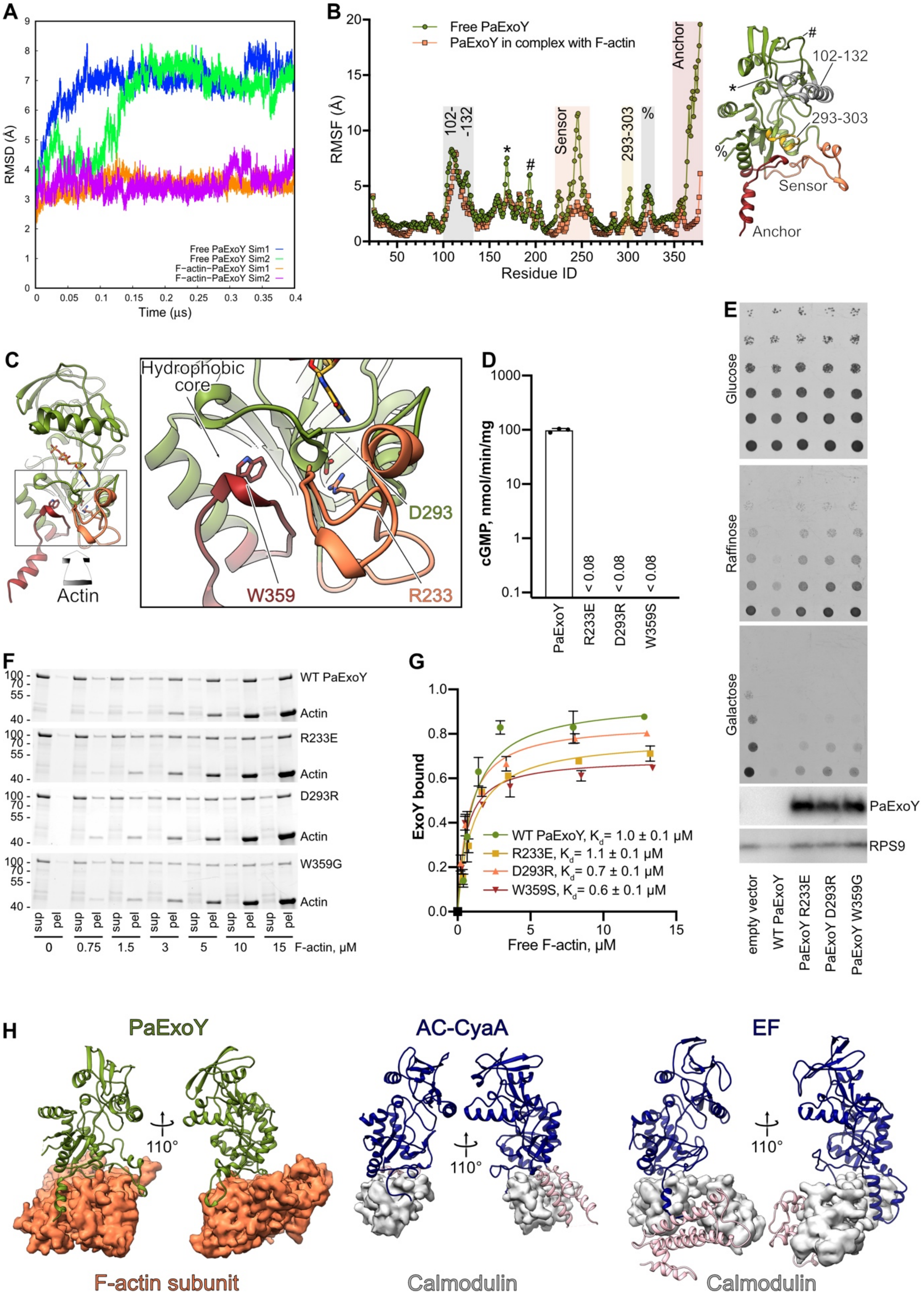

**Figure S3. Mechanism of activation of PaExoY.** (A) Root mean square fluctuation of Cα atoms of PaExoY during the simulations. The initial structure is the reference for all simulations. For each system, both independent simulations are shown separately. (B) Root mean square fluctuation of Cα atoms of ExoY. For each system, we pooled both simulations, superimposed all frames and calculated an overall RMSF. (C) Communication of activation signal through PaExoY to the nucleotide-binding pocket. (D) Activity of 30 ng of WT or 1 μg of PaExoY variants in the presence of 3 μM non-stabilized actin measured during 10 min of incubation. (E) Growth phenotype assay with yeast expressing PaExoY variants in the background (Glucose), low (Raffinose) or high (Galactose) level. Analysis of protein expression was performed by Western blot of cells grown on galactose-containing media with anti-myc (PaExoY) and anti-ribosomal protein S9 (RPS9) antibodies. (F) Cosedimentation of F-actin and 2.5 μM PaExoY detected by SDS-PAGE. Representative stain-free gels are shown. (G) The fractions of PaExoY that cosedimented with F-actin were quantified by densitometry and plotted against F-actin concentrations. (H) Comparison of PaExoY and calmodulin-activated nucleotidyl cyclases (adenylyl cyclase from *B. pertussis*, AC-CyaA (3), PDB 1YRT; edema factor from *B. anthracis*, EF, PDB 1XFV (4)). The central interaction region of the AC-CyaA and the C-terminal interaction region in EF are in pink. Protective antigen binding domain of EF is hidden for the clarity of the figure. The error bars in panels D and G correspond to standard deviations of 3 independent experiments. The uncropped gels and Western blots can be found in Fig S9.

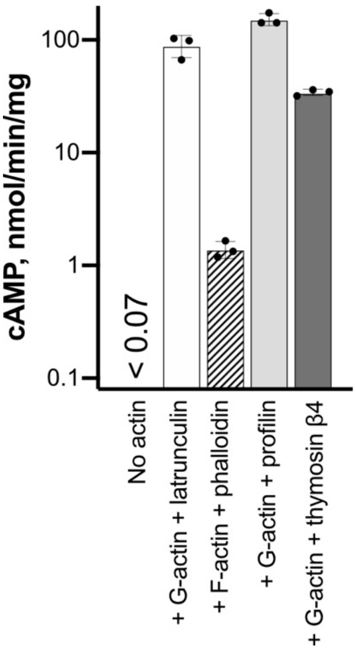

**Fig. S4. Profilin-G-actin complex activates VvExoY.** Activity of 4000 ng of VvExoY in the absence of actin; 10 ng of VvExoY in the presence of 2 μM of G-β-actin and 3 μM latrunculin A, 2 μM of G-β-actin and 3 μM profilin, or 2 μM of G-β-actin and 3 μM thymosin β4; 1000 ng of VvExoY in the presence of 2 μM of F-β-actin and 3 μM phalloidin during 10 min of incubation. The error bars correspond to standard deviations of 3 independent experiments.

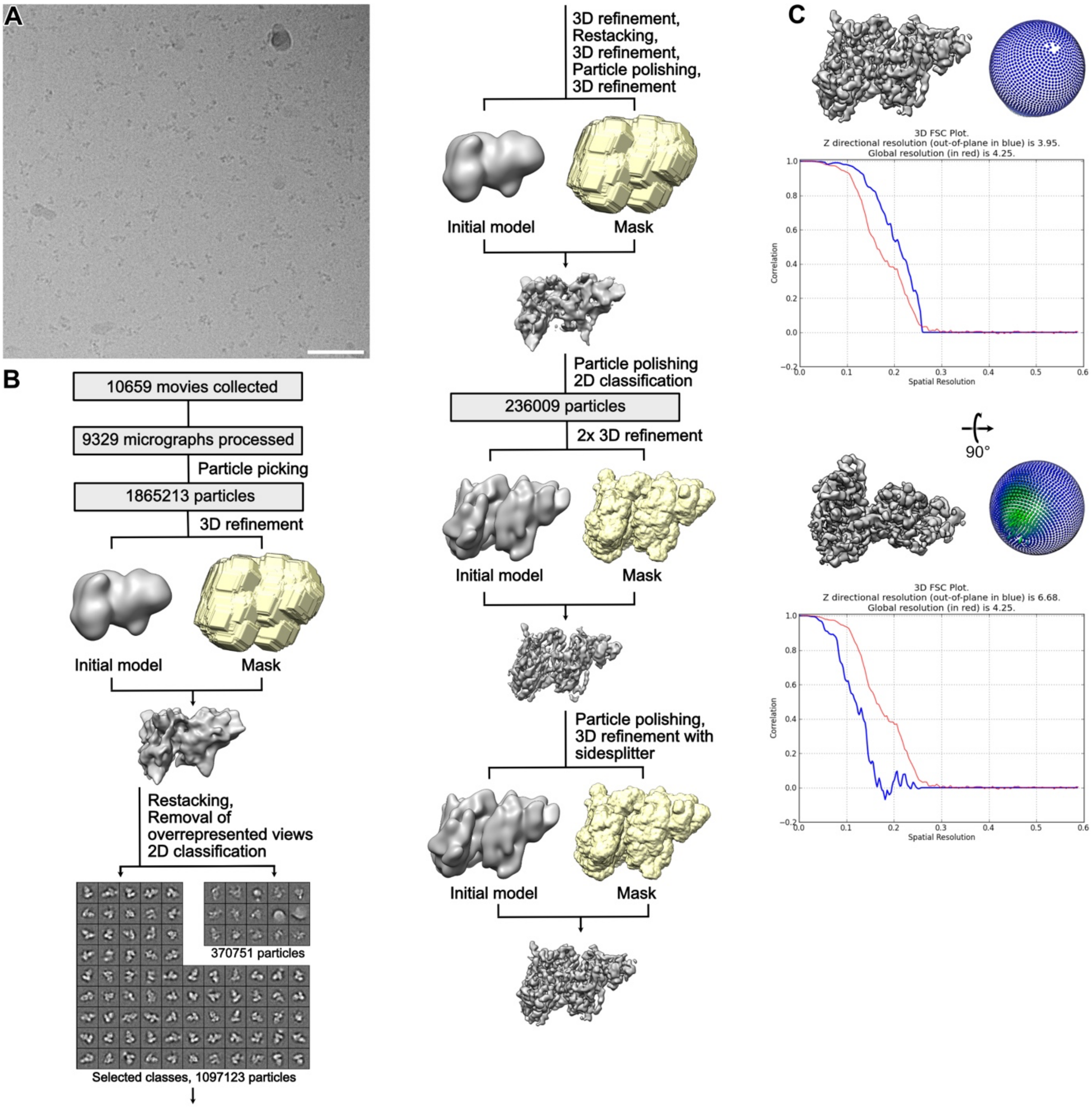

**Fig S5. Processing of the VnExoY-G-actin complex.** (A) An example cryo-EM micrograph. Scale bar 50 nm. (B) Processing overview. (C) Map postprocessed by DeemEMhancer (5), its angular distribution, and FSC plots (6) in two orientations show strong preferred specimen orientation.

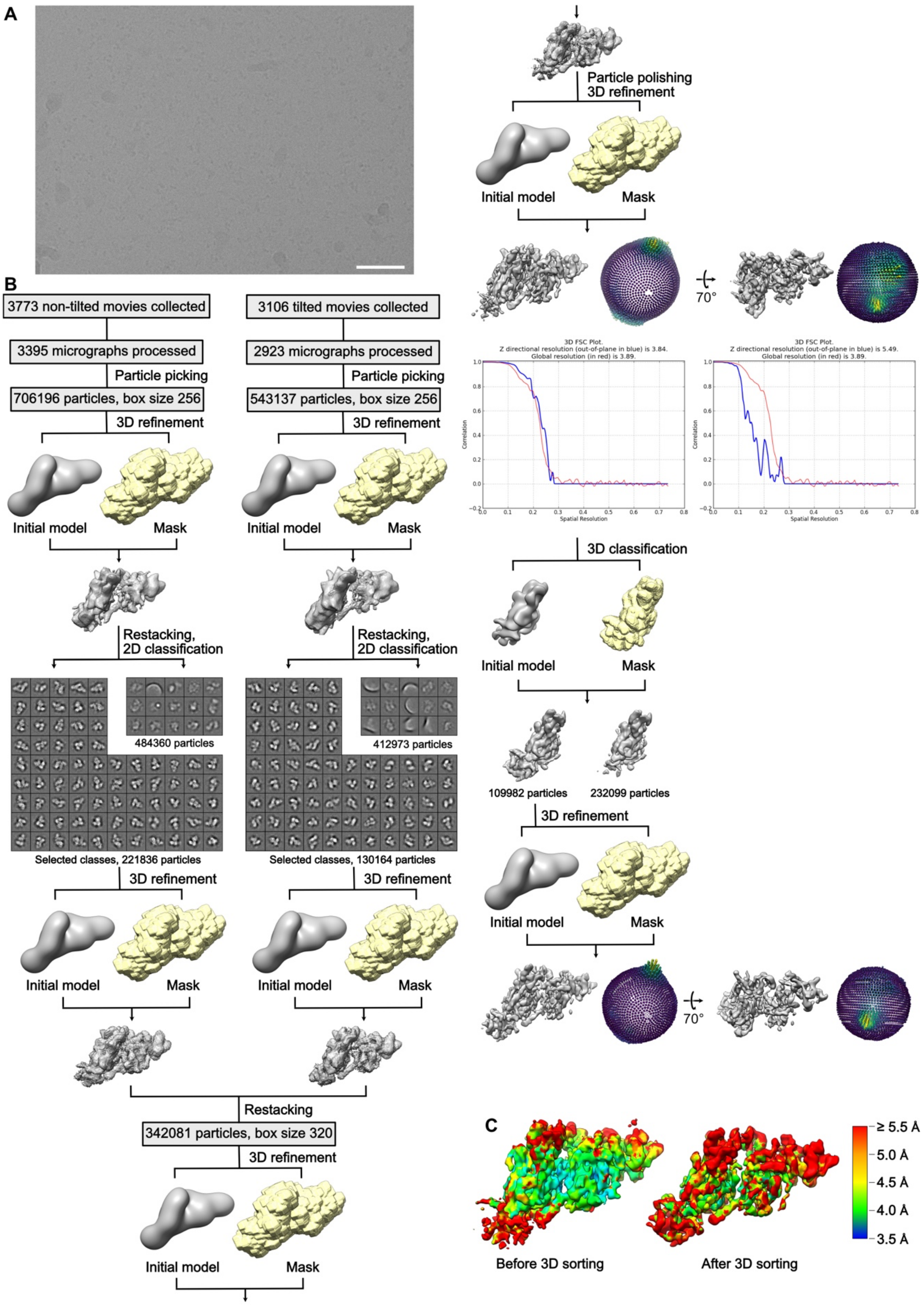

**Fig S6. Processing of the VvExoY-G-β-actin complex.** (A) An example cryo-EM micrograph. Scale bar 50 nm. (B) Processing overview with postprocessed maps, their angular distribution, and FSC plots (6). (C) Local resolution gradient of the reconstructions before and after 3D sorting. A combination of these maps was used for Fig 3A.

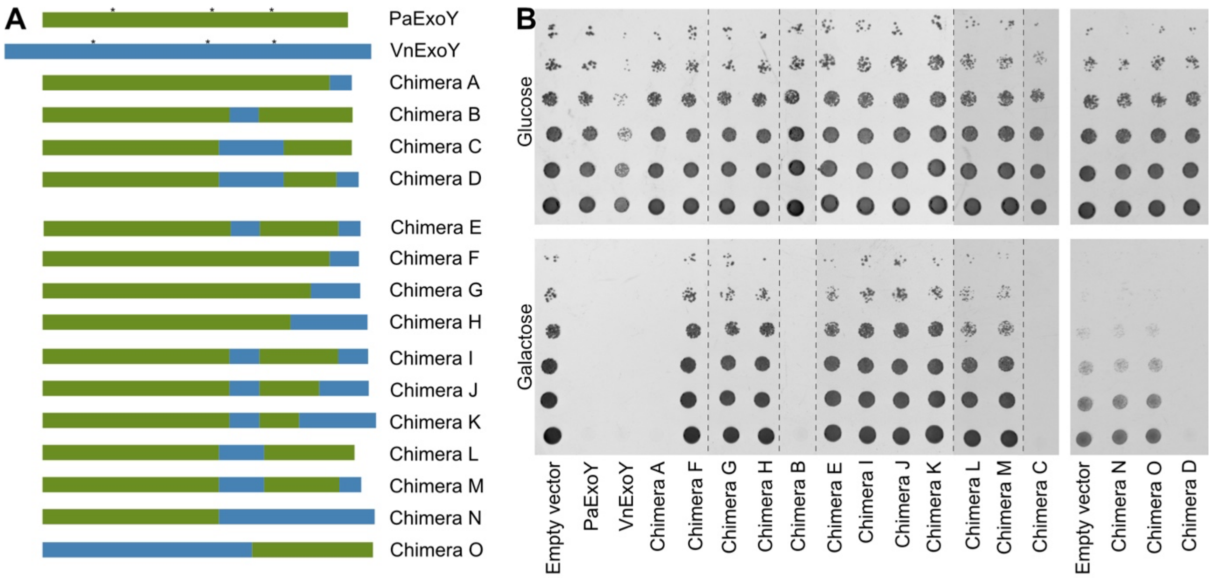

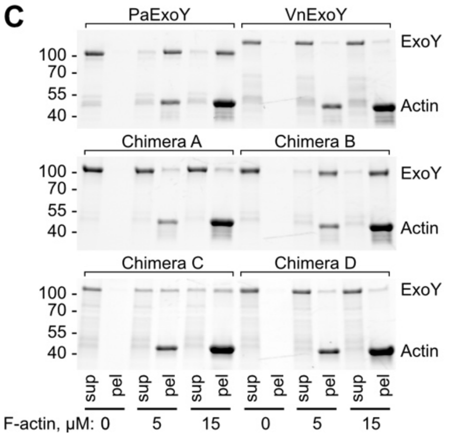

D

| | Activator | Substrate |
|---|---|---|
| PaExoY | F > G-actin | GTP > ATP |
| VnExoY | G >> F | ATP only |
| Chimera A | G = F | GTP > ATP |
| Chimera B | F > G | ATP > GTP |
| Chimera C | G > F | ATP only |
| Chimera D | G > F | ATP only |

**Fig S7. All chimera proteins created and tested in this study.** (A) A schematic representation of the created chimera proteins. (B) Yeast viability upon endogenous chimera expression on the background (Glucose) or high (Galactose) level. The panel is composed of several drop-test images as indicated by dashed lines. Amino acid sequences of the chimera proteins are available in Table S3. (C) Cosedimentation of F-actin and 2.5 μM chimera proteins detected by SDS-PAGE. Representative stain-free gels are shown. The uncropped gels can be found in Fig S9. (D) Overview of activator preference and substrate specificity of ExoY variants.

## A

F-actin activated:

```
Pa   212 DYDLFLVAPSIEAHGSGGLDARRNTAVRYTPLGAK-DPLSEDG--FYGRE-------DMARGNITPRTRQLVDALNDCLGRGEHREMFHH 291
As       DYDMFLVAPSIEEYGQGGSDARTNTAVKYQPLPN--DPHTMAS--FYGRD-------DVTKGNISPRIRDLVDTLNDTLGRGEHRDMFHH
         ***.********  * ** *** ****.* **     **        ****.        *  .***.** * *** *** *******.****

Pa   347 KGNGYHFTAHPDWNVPLRP---------SFQEALDFFQRKV  378
As       KGNGYHFNAHPDWQIPQRL---------SFQEAFAFFNQKV
         ******* *****..* *          *****. **. **
```

G-actin activated:

```
Vv   213 DYDLLTVMYSYSDLGPQDKLKQPLTWEQWKESVT-YEELTPKYKELYNSEVLYNKKDGASLGVVSDRLKALKDVINTSLGRTDGLEMVHH 301
Vn       DYDLLTVMYTYGDLGPQDKVKQPLTWEQWKESVT-YEDLSPKYKARYDNQALYEKQDGASLGMVSDRLKELKDVINTSLGRTDGLEMVHH
Va       DYDLLTVMYSYADLGPQDKVKQPLTWEQWKESVT-YEDLTPKYQELYDSEVLYNKKDGASLGVVSDRLKTLKDVINTSLGRTDGLEMVHH
Pm       DYDLFTVMYSYNDLSERSTVRKAIPWEEWRSSVN-YDELSPTYQEYYNNKDLYDRYEGEQLGIISQHVKNIKNKLNRLLQREKGKEMIHH
Vc       DYDLFSVMTHISDYGAPDMFKMPEPWADWKKSIKDPEKLDPKIKELFENEDLYNKVNNMHLGIISDRVIGLKDKINDRLGRNNGMELVHH
Vs       DYDLFSVMAHISDYGVSDLVKVPLPHAKWKATIENPDKLSPNLKELYDNEDLYNKVHNMHLGAISERVNSLKDKVNNRLGRGKGMELIHH
Vo       DYDLFTVMYSYAELGPQDKVKQPLTWEQWKESVT-YEDLTPKYKELYDSEVLYNKKDGASLGVVSDRLKTLKDVINTSLGRTDGLEMVHH
Pru      DYDLFSIIFPISELEHYAKVTEMPSWAEWKSGIN-YDELTVYQKALYENEAEYNKQEGRDNGITNSKIKEIKNKLNRRLGRTDGMELIHH
Pre      DYDLFSVIYPVSELEHYVKVSEMPSWQEWKASVN-YDELTNEQKKLYHNEAEYNKREGKDNGITNAKIKEINQELNKKLERPIGLELVHH
         ****..*.. .  ..     .         **  .   . *       ..      *         *.     .   .   .*  *  *  *.*..**

Vv   367 INVSYRASLNDKWNVGLDDPLFTPKSKLSHDFLNAKEEVIKKLSGEVETNVRTTQLLTDNEGL   425
Vn       INVSLGNYNFNWGGDGRDLGAYLGDNNNFWGGRGDDVYYATGTSNLFTGGEGQDMGVLMGREN
Va       INVSYRASLNDKWNAGLDDQLFTPKRKLSHEFLVARDKIAKMLSNESSDLGNDSELEGIAERR
Pm       INNGFISPLNEKWNEGIEGNYFSRRRKSSENYLNNRELLEKIFSDKNKYKSPLNKINNEYSDY
Vc       TNVHFTGALNTKWSENYAQL--ARKNRLSHAYLNAREQVDQLLFGKIDAAKIEFYRPKPFDET
Vs       TNVQFTGPLNSLWSKEYGQL--ARKNRLSHAYLNAREQVEALLTQGIDVAKLKVYIPGVFDEM
Vo       INVSYRASLNDKWNAGLDDPLFTPKRKLSHEFLNVRDELANKLKEKAVDFEKEVQLDRLGDHQ
Pru      INQGYRAPLNKIWNEGDNGQYFDPKRKISESFIEGRIEIARKKSLANELNDGFGYEGIRKSNR
Pre      INEGYRAPLNANWREGEYAEYFNPKRKISDSYINASREIQRKKSIPIAVRNFIEGKETQSGYI
          *  .                ..   *
```

## B

```
Pa   212 DYDLFLVAPSIEAHGSGGLDARRNTAVRYTPLGAK-DPLSEDG--FYGRE------------------DMARGNITPRTRQLVDALNDCLGRGEH-REMFHH 291
Vv   213 DYDLLTVMYSYSDLGPQDKLKQPLTWEQWKESVT-YEELTPKYKELYNSEVL-----------YNKKDGASLGVVSDRLKALKDVINTSLGRTDG-LEMVHH 301
Bps      DYDLFLIGPHITDLGPQDNLPVPDVSHAVFKARLDKYKNG------IP--S-ALRSAYEQSGEFYRKEDEEIGNASLRVRNMIPLINTALVG-DG-EAVVHH
Mt       DYDLMIVAPRAEDFGSQDIPHNPD-VSADAFAEKRPY-LL------R--RG-STKEIIRRELL--EREDPHFGNNTPRITALILQINAVLFRGEG-REVVHH
Ya       DYDLLMVAPHISDLGSQDNLPVPDVAHHVFRRRIDKY-SK------WPDAP-EIRQDYDKPASFYRKEDGEIGNASERVRQMIPYLNEAVVG-DG-EKVIHH
Pf       DYDLHMVAPHISDLGPQDRLPVPDIAHSVFTTRVDHY-RQQQPDPKAFLVPEALRADYESAEHFYQKENPDLGNATPRIEQMIRLINDRLVTPPSEERVVHH
Prs      DYDLLIGPHMRDFGHQDMLPVPDVAHSVYRQRVEKY-QR------LPTDA-NLSQAYRDENHFYQNEDSEIGNASPRVREMISLINRALVG-EA-EKVVHH
CHd      DYDLFMIAPSVTDFGFQDILPVPKDVSHSIFRKRVKQYIKD------LS--E-PLKELYSDPDTFYAPEDEEIGNASLRIRDMISDINQHLVG-NG-EKVVHH
Cd       DYDLMLIAPHVRDLGREDNLQVPDVSHQVFRQRLESY-SK------LPLHP-ELRRDYEDAASFYRRSDEEMGNSSQRVRHMVGVINQALVG-KG-EPVVHH
Cv       DYDLLVIGPHLSDVGPQDNLPVPDVAHSVFRDRIDHY-RG------IPANV-GLLDAYLNPKSFYRREDAEIGNASARIRDMIPVINQALVG-DG-ERVVHH
Als      DYDLLLVAPNLSDYGAGDLAKGDP-SQSVLESHLR-------------------NFNAPKET-TQPKAPQGFISSRVKELGDDINARLDRE-L-NKLIHH
Ei       DYDLFIASPHISDFGSQDNLLIPDVAHKVFQHRVNSYHKE----LT--T-NHELMQDYSNADSFYKNEDPDIGNASARVRILIDLINATLVG-DG-ERLVHH
         ****  .       *.              .                                             *    *   .   **  .        ..**

Pa   347 KGNGYHFTAHPDWNVPLRP---------SFQEALDFFQRKV--------------------- 378
Vv   367 INVSYRASLNDKWNVGLDDPLFTPKSKLSHDFLNAKEEVIKKLSGEVETNVRTTQLLTDNEGL 425
Bps      KNAGYHVQINPMWEKEVASV--QRESFLTARGYFSAN-TRNIHPLR-----------------
Mt       ECHGFMVPHNPLWSDEIPSL--RRE---SFEATRKLFNTL-----------------------
Ya       KRHGYHMVMNPLWEPDLLQI--RNP---HFEEMRRELILKL----------------------
Pf       KDSGYHVPLNPLWESEVVSI--KRTGFTHAQARLASAL----RQ-------------------
Prs      KAEGYHIKTNPLWEKELPTI--RRPSFEYAKRRLST-V--SLNSK------------I----
CHd      KDAGYYVPSNPLWEKDITSI--RSS---SFKAAKKVLIGHCAKIIKPK---------------
Cd       KNAGYHVPLNPLWEKEVLRA--RSP---RFDDARARIEAAM---L-ARH--------------
Cv       KDSGYHLPLNPLWEKEVSGV--RRE---SFVDARRRLGG------------------------
Als      KDQAFFVPVHNEWGQDVQNI--KRG---SFQESLDYFQKV-----------------------
Ei       RSSGYYIKINPLWEKELGKI--DSQ---RFKDAKVKLDP-L---TKPS---------------
          . .    *.

Pa   347 DYDLFLVAPSIEAHGSGGLDARRNTAVRYTPLGAK-DPLSEDG--FYGRE-------------------------------------DMARGNITPRTRQLVDALNDCLGRGEH---REMFHH 291
Vv   367 DYDLLTVMYSYSDLGPQDKLKQPLTWEQWKESVT-YEELTPKYKELYNSEVL-----------YNKKD---------------------GASLGVVSDRLKALKDVINTSLGRTDG---LEMVHH 301
Ob       DYDLMAVCPRWSDY--GSTSSAQIAK-PGLVMNGM-APRLGQVFERGTNLDKVLTMESSTGRPR--PQGWQAP--------AGF-GNEEHYDMGNVTPRILRCINLLNGRMVNGNGA--FRRVHH
Rs       DYDLMSVCPTWDSY--GGTSNAVISK-PGIRFSGK-KGLQLGQQFDVGARMDKVLDMRSNTGAR-PKLGNTKVTFQGMTKRDGG-KLAEHGDMGNITPRILRCINMLNAAMGATGGKSAGRRVHH
Mv       DYDLFAIWPFVGSY--DASAYGLDHR-PLGTVKGSVT-Q-AE-------------R---------DNVENLE--------RHFTEGGQGTKLGNITPRIYLICQLVNSIIAK------QQVLWH
En       DVDPLLEAFPLSDLDLGGKDKLPLPLISDVIVRKQVETYKNRLKSQFMEDTITAEQFVMKLSHYHDLEKELLKDFFTGITADGEHFEKEDPEMGNVTQRTRNMVSHYKRELQT--Q---EDVVHH
         *.* .     .                                                                                *  .. *          .               . *

Pa   347 KGNGYHFTAHPDWNVPLRP---------SFQEALDFFQRKV----------------------  378
Vv   367 INVSYRASLNDKWNVGLDDPLFTPKSKLSHDFLNAKEEVIKKLSGEVETNVRTTQLLTDNEGL  425
Ob       QDAGYFLPRNAAWGMSIRD-A-MRA-------V------------------------------
Rs       NEAGYYVPRNWTWGMSVRD-N-AAD---MQRYF------------------------------
Mv       DEAGIKVTLSNAWAQEPTA-QKPNRLGASYAGYVPADGVRIIVPDWYNA--------------
En       MEAGYEPEKNLLWGKTRARRPSFNQALHHLEGKTLSQQQKDKQLGEEIEKYWDSGISSESNEL
              .       *
```

**Fig S8. Alignment of PaExoY and VvExoY with known ExoY-like nucleotidyl cyclases.** (A) Eleven ExoY proteins can be clearly classified into F-actin-activated (above) and G-actin-activated (below) based on their similarity in the ABD with PaExoY and VvExoY. (B) Fourteen other ExoY-like proteins differ in the activator-binding domain from the corresponding parts in PaExoY and VvExoY and from calmodulin-binding nucleotidyl cyclases, suggesting the existence of at least one additional subgroup of bacterial nucleotidyl cyclases. Amino acids that are involved in direct contacts with actin or that organize a structural scaffold for actin-binding regions are in **bold**. Pa – *P. aeruginosa* ExoY WP_003115517, As - *Aeromonas salmonicida* ALK43954.1, Vv – *V. vulnificus* ExoY WP_039507922, Vn – *V. nigripulchritudo* WP_013610353.1, Va – *V. anguillarum* YP_004566017.1, Pm – *Proteus mirabilis* WP_020945177.1, Vc – *V. cholerae* AAW80256.1, Vs – *V. scophthalmi* 005594994.1, Vo – *V. ordalii* WP_010319615.1, Pru – *Providencia rustigianii* WP_006813565.1, Pre – *Providencia rettgeri* WP_004909604.1, Bps – *Burkholderia pseudomallei* KGC96437.1, Mt – *Mycobacterium tuberculosis* SGC81937.1, Ya – *Yersinia aldovae* WP_004701884.1, Pf – *P. fluorescens* WP_012722909.1, Prs - *Providencia* stuartii 014658369.1, CHd - *Candidatus Hamiltonella defensa* WP_015873608.1, Cd – *Cedecea davisae* WP_016538267.1, Cv – *Chromobacterium vaccinii* WP_083340618.1, Als – Algicola sagamiensis WP_083938267.1, Ei – *Edwardsiella ictaluri* YP_002932933.1, Ob – *Oxalobacteraceae* bacterium WP_020701664.1, Rs – *Rhodoferax saidenbachensis* WP_029708193.1, Mv – *Methylomerinum vadi* WP_031434121.1, En – *Endozoicomonas numazuensis* WP_034835431.1. The alignment was performed using T-Coffee (7) and adjusted manually.

Figure S2D and S3F

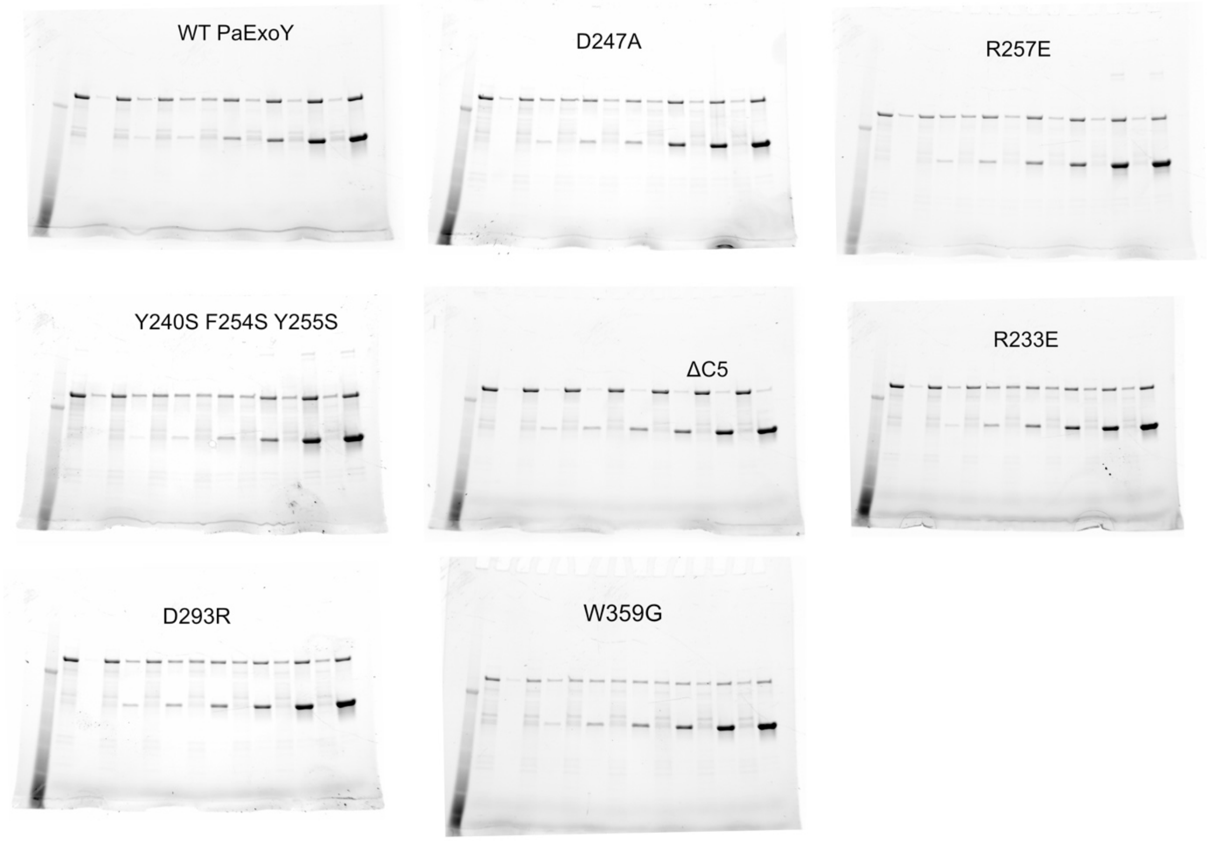

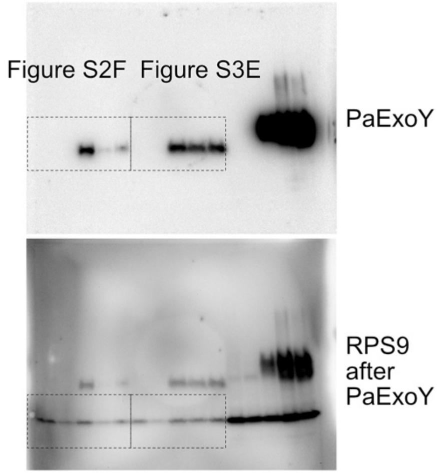

Figure S9C

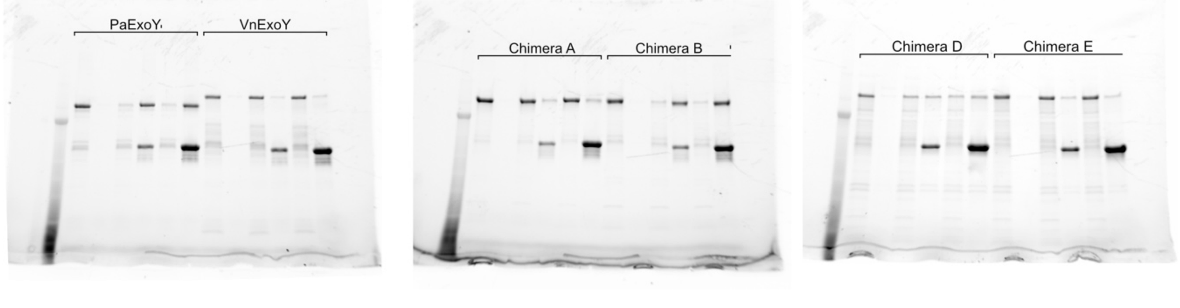

**Fig S9. Uncropped gels and Western blots.**

**Table S1. Cryo-EM data collection, refinement, and validation statistics**

| Project | PaExoY-F-α-rabbit actin | VnExoY-G-α-rabbit actin | VvExoY-G-β-human actin-profilin |
|---|---|---|---|
| Microscope | Titan Krios | Titan Krios | Titan Krios |
| Voltage (kV) | 300 | 300 | 300 |
| Defocus range (µm) | -0.4 to -3 | -1 to -2.5 | -1.2 to -2.5 |
| Camera | Falcon III (Linear mode) | Gatan K2 (Counting mode) | Gatan K3 (Superresolution mode) |
| Pixel size (Å) | 1.1 | 0.85 | 0.34, 0.68 binned |
| Total electron dose (e/Å$^2$) | 93 | 80 | 60 |
| Exposure time (s) | 1.5 | 8 | 2 |
| Frames per movie | 40 | 64 | 60 |
| Number of movies | 8,663 (12,437) | 9,329 (10,659) | 6,318 (6,880) |
| **3D Refinement** | | | |
| Number of particles | 1,535,755 (2,249,589) | 236,009 (1,865,213) | 342,081 (1,299,405) |
| Final resolution (Å) | 3.2 | 4.2 | 3.9 |
| Helical rise (Å) | 27 | - | - |
| Helical twist (°) | -166.9 | - | - |
| **Atomic model statistics** | | | |
| Non-hydrogen atoms | 28585 | - | 7073 |
| Molprobity score | 1.95 | - | 1.98 |
| Clashscore | 10.58 | - | 12.09 |
| EMRinger score | 3.43 | - | 1.49 |
| Bond RMSD (Å) | 0.021 | - | 0.005 |
| Angle RMSD (°) | 1.471 | - | 1.024 |
| Poor rotamers (%) | 0.17 | - | 0 |
| Favored rotamers (%) | 95.13 | - | 92.95 |
| Ramachandran favored (%) | 93.36 | - | 94.25 |
| Ramachandran allowed (%) | 6.32 | - | 5.75 |
| Ramachandran outliers (%) | 0 | - | 0 |

**Table S2**. **List of primers, strains and plasmids used in this study.**

| **Bacterial and yeast strains** | **Description** | **Reference** |
|---|---|---|
| *E. coli* DH5α | F⁻ Φ80*lac*ZΔM15 Δ(*lac*ZYA-*arg*F) U169 *rec*A1 *end*A1 *hsd*R17($r_k^-$, $m_k^+$) *pho*A *sup*E44 *thi*-1 *gyr*A96 *rel*A1 λ⁻ | Invitrogen |
| *E. coli* BL21 DE3 CodonPlus RIPL | F⁻ *ompT hsdS*($r_B^-$ $m_B^-$) *dcm*⁺ Tet$^r$ *gal*λ(DE3) *endA* Hte [*argU proL* Cam$^r$] [*argU ileY leuW* Strep/Spec$^r$] | Agilent |
| *S. cerevisiae* MH272-3fα | "Wild-type" strain, *ura3, leu2, his3, trp1, ade2* | (8) |
| *S. cerevisiae* Y395 | *S. cerevisiae* MH272-3fα + Vector[Ade] (2473) | (2) |
| *S. cerevisiae* Y410 | *S. cerevisiae* MH272-3fα + PaExoY[Ade] (p1593) | (2) |
| *S. cerevisiae* Y655 | *S. cerevisiae* MH272-3fα + VnExoY[Ade] (p1648) | (9) |
| *S. cerevisiae* Y642 | *S. cerevisiae* MH272-3fα + PaExoY_Y240S_F254S_Y255S[Ade] (pB636) | This study |
| *S. cerevisiae* Y595 | *S. cerevisiae* MH272-3fα + PaExoY_D247A[Ade] (pB568) | This study |
| *S. cerevisiae* Y593 | *S. cerevisiae* MH272-3fα + PaExoY_R257E[Ade] (pB566) | This study |
| *S. cerevisiae* Y532 | *S. cerevisiae* MH272-3fα + PaExoY_R233E[Ade] (pB490) | This study |
| *S. cerevisiae* Y603 | *S. cerevisiae* MH272-3fα + PaExoY_D293R[Ade] (pB582) | This study |
| *S. cerevisiae* Y608 | *S. cerevisiae* MH272-3fα + PaExoY_W359S[Ade] (pB587) | This study |
| *S. cerevisiae* Y722 | *S. cerevisiae* MH272-3fα + VnExoY_W363S[Ade]( pB737) | This study |
| *S. cerevisiae* Y728 | *S. cerevisiae* MH272-3fα + VnExoY_W222S_W225S_Y248S[Ade] (pB743) | This study |
| *S. cerevisiae* Y732 | *S. cerevisiae* MH272-3fα + VnExoY_ΔC27[Ade] (pB747) | This study |
| *S. cerevisiae* Y656 | *S. cerevisiae* MH272-3fα + ExoY_chimeraA[Ade] (pB672) | This study |
| *S. cerevisiae* Y664 | *S. cerevisiae* MH272-3fα + ExoY_chimeraB[Ade] (pB678) | This study |
| *S. cerevisiae* Y665 | *S. cerevisiae* MH272-3fα + ExoY_chimeraC[Ade] (pB681) | This study |
| *S. cerevisiae* Y747 | *S. cerevisiae* MH272-3fα + ExoY_chimeraD[Ade] (pB756) | This study |
| *S. cerevisiae* Y717 | *S. cerevisiae* MH272-3fα + ExoY_chimeraE[Ade] (pB731) | This study |
| *S. cerevisiae* Y657 | *S. cerevisiae* MH272-3fα + ExoY_chimeraF[Ade] (pB673) | This study |
| *S. cerevisiae* Y660 | *S. cerevisiae* MH272-3fα + ExoY_chimeraG[Ade] (pB679) | This study |
| *S. cerevisiae* Y661 | *S. cerevisiae* MH272-3fα + ExoY_chimeraH[Ade] (pB680) | This study |
| *S. cerevisiae* Y666 | *S. cerevisiae* MH272-3fα + ExoY_chimeraI[Ade] (pB682) | This study |
| *S. cerevisiae* Y667 | *S. cerevisiae* MH272-3fα + ExoY_chimeraJ[Ade] (pB683) | This study |
| *S. cerevisiae* Y668 | *S. cerevisiae* MH272-3fα + ExoY_chimeraL[Ade] (pB684) | This study |
| *S. cerevisiae* Y697 | *S. cerevisiae* MH272-3fα + ExoY_chimeraL[Ade] (pB717) | This study |
| *S. cerevisiae* Y698 | *S. cerevisiae* MH272-3fα + ExoY_chimeraM[Ade] (pB719) | This study |
| *S. cerevisiae* Y715 | *S. cerevisiae* MH272-3fα + ExoY_chimeraN[Ade] (pB729) | This study |
| *S. cerevisiae* Y716 | *S. cerevisiae* MH272-3fα + ExoY_chimeraO[Ade] (pB730) | This study |
| **Plasmids for experiments in *S. cerevisiae*** | | |
| 2473 YEpGal555 | *E. coli*/*S. cerevisiae* shuttle vector [ADE2] with Gal1 promoter | (10) |
| p1593 YEpGal555 PaExoY | WT PaExoY with N-terminal myc-tag in YEpGal555 vector | (2) |
| p1648 YEpGal555 VnExoY | WT VnExoY with N-terminal myc-tag in YEpGal555 vector | (9) |
| pB576 YEpGal555 PaExoY_Y240S | The Y240S mutation was generated by two-step overlap PCR using oligonucleotides cagcgaagcgatgatttttg, caggttgtctaactccttcc, | This study |

| | | |
|---|---|---|
| | ccgcggtcagatccacccccctcgg and ccgagggggggtggatctgaccgcgg, and p1593 as a matrix. The PCR product was digested with XhoI and KpnI and ligated into digested 2473 YEpGal555 vector. | |
| pB636 YEpGal555 PaExoY_Y240S_F254S_Y255S | The F254S and Y255S mutations were generated by two-step overlap PCR using oligonucleotides cagcgaagcgatgatttttg, caggttgtctaactccttcc, ctgagcgaggacggatccagtggcagggaggatatg and catatcctccctgccactggatccgtcctcgctcag, and pB576 as a matrix. The PCR product was digested with XhoI and KpnI and ligated into digested 2473 YEpGal555 vector. | This study |
| pB568 YEpGal555 PaExoY_D247A | The D247A mutation was generated by two-step overlap PCR using oligonucleotides cagcgaagcgatgatttttg, caggttgtctaactccttcc, cggtgcaaagcttcccctgagcgag and ctcgctcaggggaagctttgcaccg, and p1593 as a matrix. The PCR product was digested with XhoI and KpnI and ligated into digested 2473 YEpGal555 vector. | This study |
| pB566 YEpGal555 PaExoY_D257E | The D257E mutation was generated by two-step overlap PCR using oligonucleotides cagcgaagcgatgatttttg, caggttgtctaactccttcc, ggattctatggcgaggaggatatggcc and ggccatatcctcctcgccatagaatcc, and p1593 as a matrix. The PCR product was digested with XhoI and KpnI and ligated into digested 2473 YEpGal555 vector. | This study |
| pB490 YEpGal555 PaExoY_R233E | The R233E mutation was generated by two-step overlap PCR using oligonucleotides cagcgaagcgatgatttttg, caggttgtctaactccttcc, ggtctcgatgcagaaaggaataccg and cggtattcctttctgcatcgagacc, and p1593 as a matrix. The PCR product was digested with XhoI and KpnI and ligated into digested 2473 YEpGal555 vector. | This study |
| pB582 YEpGal555 PaExoY_D293R | The D293R mutation was generated by two-step overlap PCR using oligonucleotides cagcgaagcgatgatttttg, caggttgtctaactccttcc, gtttcaccacagccgcgatgcgggcaaccc and gggttgcccgcatcgcggctgtggtgaaac, and p1593 as a matrix. The PCR product was digested with XhoI and KpnI and ligated into digested 2473 YEpGal555 vector. | This study |
| pB587 YEpGal555 PaExoY_W359S | The W359S mutation was generated by two-step overlap PCR using oligonucleotides cagcgaagcgatgatttttg, caggttgtctaactccttcc, gcccatcccgactcgaacgtgccgc and gcggcacgttcgagtcgggatgggc, and p1593 as a matrix. The PCR product was digested with XhoI and KpnI and ligated into digested 2473 YEpGal555 vector. | This study |
| pB737 YEpGal555 VnExoY_W363S | The W363S mutation was generated by two-step overlap PCR using oligonucleotides cagcgaagcgatgatttttg, caggttgtctaactccttcc, ctcaatgataagtcgaattcgggctta and taagcccgaattcgacttatcattgag,and p1648 as a matrix. The PCR product was digested with XhoI and KpnI and ligated into digested 2473 YEpGal555 vector. | This study |
| pB742 YEpGal555 VnExoY_Y248S | The Y248S mutation was generated by two-step overlap PCR using oligonucleotides cagcgaagcgatgatttttg, caggttgtctaactccttcc, caatcaagctctttcggagaaacaggatg and catcctgtttctccgaaagagcttgattg,and p1648 as a matrix. The PCR product was digested with XhoI and KpnI and ligated into digested 2473 YEpGal555 vector. | This study |
| pB743 YEpGal555 VnExoY_ W222S_W225S_Y248S | The W222S and W225S mutations were generated by two-step overlap PCR using oligonucleotides cagcgaagcgatgatttttg, caggttgtctaactccttcc, gcaaccgctcacgtcggaacagtcgaaagaatcgg and | This study |

| | | |
|---|---|---|
| | ccgattctttcgactgttccgacgtgagcggttgc,and pB742 as a matrix. The PCR product was digested with XhoI and KpnI and ligated into digested 2473 YEpGal555 vector. | |
| pB747 YEpGal555 VnExoY_ΔC27 | C-terminal deletion was generated by a PCR using oligonucleotides cagcgaagcgatgatttttg and tataggtacctagcttacgttttgtagtgaac, and pB 1648 as a matrix. The PCR product was digested with XhoI and KpnI and ligated into digested 2473 YEpGal555 vector. | This study |
| pB672 YEpGal555 ChimeraA | The C-terminus of PaExoY was substituted by the homological region of VnExoY by a PCR using oligonucleotides agacctcgagcgtatcgacggtcatcgtcagg and tataggtaccgagtccgttgagcttcgaagattctgtcaaacccagctttttcgccacttcatcccgagcgtctaagtaatcgggccgcagcggcacgttccagtcggg, and p1593 as a matrix. The PCR product was digested with XhoI and KpnI and ligated into digested 2473 YEpGal555 vector. | This study |
| pB678 YEpGal555 ChimeraB | The central region of PaExoY was substituted by the homological region of VnExoY by the two-step overlap PCR using oligonucleotides cagcgaagcgatgatttttg, caggttgtctaactccttcc, gacttatcacctaagtataaagcgcgttatgacaatcaagctctttacgagaaacaggatggcgcaagtttgggaaacatcactccgcgcacgcggcaac and cgcgctttatacttaggtgataagtcttcgtaggttaccgattctttccactgttcccacgtgagcggttgccttcttgcatcgagaccaccactgccatg. The PCR product was digested with XhoI and KpnI and ligated into digested 2473 YEpGal555 vector. | This study |
| pB681 YEpGal555 ChimeraE | The C-terminus of PaExoY was substituted by the homological region of VnExoY by a PCR using oligonucleotides agacctcgagcgtatcgacggtcatcgtcagg and tataggtaccgagtccgttgagcttcgaagattctgtcaaacccagctttttcgccacttcatcccgagcgtctaagtaatcgggccgcagcggcacgttccagtcggg, and pB678 as a matrix. The PCR product was digested with XhoI and KpnI and ligated into digested 2473 YEpGal555 vector. | This study |
| pB756 YEpGal555 ChimeraC | The C-terminus of PaExoY was substituted by the homological region of VnExoY by swaping NcoI/KpnI fragment of pB731 with the similar fragment of p1593. | This study |
| pB731 YEpGal555 ChimeraD | The chimera gene was synthesized by Twist Bioscience, digested with XhoI and KpnI and ligated into digested 2473 YEpGal555 vector. | This study |
| pB673 YEpGal555 ChimeraF | The C-terminus of VnExoY was first PCR-amplified using oligonucleotides gccgctgcggcccctgttcactacaaaacgtaagctatc and caggttgtctaactccttcc from the matrix p1648. The remaining part of the toxin was PCR-amplified using oligonucleotides ttgtagtgaacaggggccgcagcggcacgttccag and cagcgaagcgatgatttttg from the plasmid p1593. After the PCR reaction with oligonucleotides cagcgaagcgatgatttttg and caggttgtctaactccttcc, and the products of the previous reactions as matrixes, the complete chimera gene was digested with XhoI and KpnI and ligated into digested 2473 YEpGal555 vector. | This study |
| pB679 YEpGal555 ChimeraG | The C-terminus of VnExoY was first PCR-amplified using oligonucleotides cggaagagcttttccaattttcagcaggtcgcgattaatgcg and caggttgtctaactccttcc from the matrix p1648. The remaining part of the toxin was PCR-amplified using oligonucleotides cgacctgctgaaaattggaaaagctcttccgatccgccacc and cagcgaagcgatgatttttg from the plasmid p1593. After the PCR reaction with oligonucleotides cagcgaagcgatgatttttg and caggttgtctaactccttcc, and the products of the previous reactions as matrixes, the complete chimera gene was digested with XhoI and KpnI and ligated into digested 2473 YEpGal555 vector. | This study |
| pB680 YEpGal555 ChimeraH | The C-terminus of VnExoY was first PCR-amplified using oligonucleotides ccttccccggcactttttgatgatgatggtctgg and caggttgtctaactccttcc from the matrix p1648. The remaining part of the toxin was PCR-amplified using oligonucleotides | This study |

| | | |
|---|---|---|
| | catcatcaaaaaagtgccggggaaggtagaaggtggccg and cagcgaagcgatgatttttg from the plasmid p1593. After the PCR reaction with oligonucleotides cagcgaagcgatgatttttg and caggttgtctaactccttcc, and the products of the previous reactions as matrixes, the complete chimera gene was digested with XhoI and KpnI and ligated into digested 2473 YEpGal555 vector. | |
| pB682 YEpGal555 ChimeraI | The C-terminus of VnExoY was introduced into pB678 by digestion of pB673 with NdeI and exchanging of the segment with the similar region of NdeI-digested pB678 | This study |
| pB683 YEpGal555 ChimeraJ | The C-terminus of VnExoY was introduced into pB678 by digestion of pB679 with NdeI and exchanging of the segment with the similar region of NdeI-digested pB678 | This study |
| pB684 YEpGal555 ChimeraK | The C-terminus of VnExoY was introduced into pB678 by digestion of pB680 with NdeI and exchanging of the segment with the similar region of NdeI-digested pB678 | This study |
| pB717 YEpGal555 ChimeraL | The central region of PaExoY was substituted by the homological region of VnExoY by the two-step overlap PCR using oligonucleotides cagcgaagcgatgatttttg, caggttgtctaactccttcc, gacctcttcctggtagcgcccacctatggggatttaggtccacaggataaggtgaagcaaccgctcacgtgggaacagtggaaagaatcggtaacctacgaagacttatcacctaagtat and tccaccagttgccgcgtgcgtttcaagcgatcgctgaccatgcccaaacttgcgccatcctgtttctcgtaaagagcttgattgtcataacgcgctttatacttaggtgataagtcttcg, and p1593 as a matrix. The PCR product was digested with XhoI and KpnI and ligated into digested 2473 YEpGal555 vector. | This study |
| pB719 YEpGal555 ChimeraM | The C-terminus of PaExoY was substituted by the homological region of VnExoY by a PCR using oligonucleotides agacctcgagcgtatcgacggtcatcgtcagg and tataggtaccgagtccgttgagcttcgaagattctgtcaaacccagctttttcgccacttcatcccgagcgtctaagtaatcgggccgcagcggcacgttccagtcggg, and pB717 as a matrix. The PCR product was digested with XhoI and KpnI and ligated into digested 2473 YEpGal555 vector. | This study |
| pB729 YEpGal555 ChimeraN | The chimera was generated using two-step overlap PCR. 5'of the chimera was PCR-amplified using oligonucleotides cagcgaagcgatgatttttg and cctaaatccccataggtgggcgctaccaggaagaggtcataatc, and matrix p1593. The 3'of the chimera was PCR-amplified using oligonucleotides caggttgtctaactccttcc and cttcctggtagcgcccacctatggggatttaggtccacagg, and matrix p1648. The PCR-fragments of the second PCR step with oligonucleotides caggttgtctaactccttcc and cagcgaagcgatgatttttg, and PCR-fragments from the previous reactions, was digested with XhoI and KpnI and ligated into digested 2473 YEpGal555 vector. | This study |
| pB730 YEpGal555 ChimeraO | The chimera was generated using two-step overlap PCR. 5'of the chimera was PCR-amplified using oligonucleotides cagcgaagcgatgatttttg and catgcgcctcgatcgagtacatcaccgttaacaagtcataatcc, and matrix p1648. The 3'of the chimera was PCR-amplified using oligonucleotides caggttgtctaactccttcc and gttaacggtgatgtactcgatcgaggcgcatggcagtggtg, and matrix p1593. The PCR-fragments of the second PCR step with oligonucleotides caggttgtctaactccttcc and cagcgaagcgatgatttttg, and PCR-fragments from the previous reactions, was digested with XhoI and KpnI and ligated into digested 2473 YEpGal555 vector. | This study |
| **Plasmids for protein expression in *E. coli*** | | |
| 2479 pB386 pET28a MBP-His-ExoY | WT MBP-PaExoY fusion protein with N-terminal His-tag | (11) |

| | | |
|---|---|---|
| pUM522 | WT VnExoY with C-terminal His-tag | (12) |
| pB642 pET28a MBP-PaExoY D247A | ExoY gene with the mutation was amplified from pB568 using oligonucleotides tatagagctctggtcgtatcgacggtcatcgtca and tataaagcttcagaccttacgttggaaaaagtc, digested with SacI and HindIII and ligated into digested pB137 vector (2) | This study |
| pB643 pET28a MBP-PaExoY R257E | ExoY gene with the mutation was amplified from pB566 using oligonucleotides tatagagctctggtcgtatcgacggtcatcgtca and tataaagcttcagaccttacgttggaaaaagtc, digested with SacI and HindIII and ligated into digested pB137 vector (2) | This study |
| pB644 pET28a MBP-PaExoY Y240A F254S Y255S | ExoY gene with the mutation was amplified from pB636 using oligonucleotides tatagagctctggtcgtatcgacggtcatcgtca and tataaagcttcagaccttacgttggaaaaagtc, digested with SacI and HindIII and ligated into digested pB137 vector (2) | This study |
| pB610 pET28a MBP-PaExoY R233E | ExoY gene with the mutation was amplified from pB490 using oligonucleotides tatagagctctggtcgtatcgacggtcatcgtca and tataaagcttcagaccttacgttggaaaaagtc, digested with SacI and HindIII and ligated into digested pB137 vector (2) | This study |
| pB637 pET28a MBP-PaExoY D293R | ExoY gene with the mutation was amplified from pB582 using oligonucleotides tatagagctctggtcgtatcgacggtcatcgtca and tataaagcttcagaccttacgttggaaaaagtc, digested with SacI and HindIII and ligated into digested pB137 vector (2) | This study |
| pB638 pET28a MBP-PaExoY W359S | ExoY gene with the mutation was amplified from pB587 using oligonucleotides tatagagctctggtcgtatcgacggtcatcgtca and tataaagcttcagaccttacgttggaaaaagtc, digested with SacI and HindIII and ligated into digested pB137 vector (2) | This study |
| pB575 pET28a MBP-PaExoY ΔC5 | C-terminal deletion was generated by PCR reaction with oligonucleotides tatagagctctggtcgtatcgacggtcatcgtca and tatacaagcttcaaaagtcgagcgcctcctggaag, and 2479 pB386 pET28a MBP-His-ExoY as a matrix. The amplified gene was digested with SacI and HindIII and ligated into digested pB137 vector (2) | This study |
| pB686 pET28a MBP | SalI and KpnI digesting sites were introduced into pB137 vector (2) using oligonucleotides ctgtcgacggttgataaggtacctaagctagctaaa and agcttttagctagcttaggtaccttatcaaccgtcgacagagct to simplify cloning of genes with N-terminal MBP tag | This study |
| pB687 pET28a MBP-VnExoY | XhoI KpnI fragment of p1648 was ligated into digested with SalI and KpnI pB686 plasmid to generate a fusion protein of VnExoY and MBP | This study |
| pB769 MBP-VnExoY W363S | XhoI KpnI fragment of pB737 was ligated into digested with SalI and KpnI pB686 to generate a fusion protein of VnExoY with the mutation and MBP | This study |
| pB770 MBP-VnExoY W222S W225S Y248S | XhoI KpnI fragment of pB743 was ligated into digested with SalI and KpnI pB686 to generate a fusion protein of VnExoY with the mutation and MBP | This study |
| pB771 MBP-PaExoY ΔC27 | XhoI KpnI fragment of pB747 was ligated into digested with SalI and KpnI pB686 to generate a fusion protein of VnExoY with the mutation and MBP | This study |
| pB693 MBP-VvExoY | VvExoY gene was synthesized by Twist Bioscience, digested with XhoI and KpnI and ligated into SalI/KpnI digested pB686. | This study |
| pB690 pET28a MBP-Chimera A | XhoI KpnI fragment of pB672 was ligated into digested with SalI and KpnI pB686 to generate a fusion protein of the chimera and MBP | This study |
| pB691 pET28a MBP-Chimera B | XhoI KpnI fragment of pB678 was ligated into digested with SalI and KpnI pB686 to generate a fusion protein of the chimera and MBP | This study |
| pB775 pET28a MBP-Chimera C | XhoI KpnI fragment of pB756 was ligated into digested with SalI and KpnI pB686 to generate a fusion protein of the chimera and MBP | This study |
| pB733 pET28a MBP-Chimera D | XhoI KpnI fragment of pB731 was ligated into digested with SalI and KpnI pB686 to generate a fusion protein of the chimera and MBP | This study |
| **Plasmids for protein expression in insect cells** | | |

| | | |
|---|---|---|
| p2098 pFL_ACTB | First, ActB gene was amplified with oligonucleotides tatataggatccatggatgatgatatcgccgcgctc and tatataaagcttctagaagcatttgcggtggacgatg from cDNA clone (BioCat), digested with BamHI and HindIII, and inserted into pFL vector. Then, the nucleotide sequence encoding a cleavable linker and thymosin β4 were introduced downstream of actin gene using Gibson cloning. | This study |
| p2336 pFL_ACTB_C272A | C272A mutation was introduced by the QuikChange method using oligonucleotides cttcctgggcatggagtccgctggcatccacgaaactaccttc and gaaggtagtttcgtggatgccagcggactccatgcccaggaag and p2098 as a matrix. | This study |

### Table S3. Amino acid sequence of PaExoY-VnExoY chimeras

Green and blue colors correspond to PaExoY and VnExoY sequence, respectively.

| ID | Plasmid | Amino acid sequence |
|---|---|---|
| WT PaExoY, p1593 | | meqkliseedleridghrqvvsnataqpgpllrpadmqaralqdlfdaqgvgvpvehalrmqavarqtntvfgirpverivttlieegfptkgfsvkgkssnwgpqagficvdqhlskredrdtaeirklnlavakgmdggaytqtdlrisrqrlaelvrnfglvadgvgpvrlltaqgpsgkryefearqepdglyrisrlgrseavqvlaspacglamtadydlflvapsieahgsggldarrntavrytplgakdplsedgfygredmargnitprtrqlvdalndclgrgehremfhhsddagnpgshmgdnfpatfylpramehrvgeesvrfdevcvvadrksfsllvecikgngyhftahpdwnvplrpsfqealdffqrkvnpgtaas* |
| WT VnExoY, p1648 | | meqkliseedlegynygqalqeaqldiatmkprqrvtanelqlgddnaitnavtseqeatpnqdgshktyqsrdlvlepiqhpksielgmpevdqsvlaevaerenviigvrpvdeksksliaskmysskglfvkakssdwgpmsgfipvdqsfakasarrdlekfneyaeqsilsgnavsanlylnqvrieelvskyesltpleldvdsgmykttatngdqtipfflnkvtvddkelwqvhylregelapfkvigdpvskqpmtadydlltvmytygdlgpqdkvkqpltweqwkesvtyedlspkykarydnqalyekqdgaslgmvsdrlkelkdvintslgrtdglemvhhgaddanpyavmadnfpatffvpkhffdddglgegkgsiqtyfnvneqgavviqnpqefsnfqqvainasyraslndkwnsgldsplfttkrklshdyldardevakklgltessklnglgtaas* |
| A | pB672 | meqkliseedleridghrqvvsnataqpgpllrpadmqaralqdlfdaqgvgvpvehalrmqavarqtntvfgirpverivttlieegfptkgfsvkgkssnwgpqagficvdqhlskredrdtaeirklnlavakgmdggaytqtdlrisrqrlaelvrnfglvadgvgpvrlltaqgpsgkryefearqepdglyrisrlgrseavqvlaspacglamtadydlflvapsieahgsggldarrntavrytplgakdplsedgfygredmargnitprtrqlvdalndclgrgehremfhhsddagnpgshmgdnfpatfylpramehrvgeesvrfdevcvvadrksfsllvecikgngyhftahpdwnvplrpdyldardevakklgltessklnglgtaas* |
| B | pB678 | meqkliseedleridghrqvvsnataqpgpllrpadmqaralqdlfdaqgvgvpvehalrmqavarqtntvfgirpverivttlieegfptkgfsvkgkssnwgpqagficvdqhlskredrdtaeirklnlavakgmdggaytqtdlrisrqrlaelvrnfglvadgvgpvrlltaqgpsgkryefearqepdglyrisrlgrseavqvlaspacglamtadydlflvapsieahgsggldarrqpltweqwkesvtyedlspkykarydnqalyekqdgaslgnitprtrqlvdalndclgrgehremfhhsddagnpgshmgdnfpatfylpramehrvgeesvrfdevcvvadrksfsllvecikgngyhftahpdwnvplrpsfqealdffqrkvnpgtaas* |
| C | pB756 | meqkliseedleridghrqvvsnataqpgpllrpadmqaralqdlfdaqgvgvpvehalrmqavarqtntvfgirpverivttlieegfptkgfsvkgkssnwgpqagficvdqhlskredrdtaeirklnlavakgmdggaytqtdlrisrqrlaelvrnfglvadgvgpvrlltaqgpsgkryefearqepdglyrisrlgrseavqvlaspacglamtadydlflvaptygdlgpqdkvkqpltweqwkesvtyedlspkykarydnqalyekqdgaslgmvsdrlkelkdvintslgrtdglemvhhgaddanpgshmgdnfpatfylpramehrvgeesvrfdevcvvadrksfsllvecikgngyhftahpdwnvplrpsfqealdffqrkvnpgtaas* |
| D | pB731 | meqkliseedleridghrqvvsnataqpgpllrpadmqaralqdlfdaqgvgvpvehalrmqavarqtntvfgirpverivttlieegfptkgfsvkgkssnwgpqagficvdqhlskredrdtaeirklnlavakgmdggaytqtdlrisrqrlaelvrnfglvadgvgpvrlltaqgpsgkryefearqepdglyrisrlgrseavqvlaspacglamtadydlflvaptygdlgpqdkvkqpltweqwkesvtyedlspkykarydnqalyekqdgaslgmvsdrlkelkdvintslgrtdglemvhhgaddanpgshmgdnfpatfylpramehrvgeesvrfdevcvvadrksfsllvecikgngyhftahpdwnvplrpdyldardevakklgltessklnglgtaas* |
| E | pB681 | meqkliseedleridghrqvvsnataqpgpllrpadmqaralqdlfdaqgvgvpvehalrmqavarqtntvfgirpverivttlieegfptkgfsvkgkssnwgpqagficvdqhlskredrdtaeirklnlavakgmdggaytqtdlrisrqrlaelvrnfglvadgvgpvrlltaqgpsgkryefearqepdglyrisrlgrseavqvlaspacglamtadydlflvapsieahgsggldarrqpltweqwkesvtyedlspkykarydnqalyekqdgaslgnitprtrqlvdalndclgrgehremfhhsddagnpgshmgdnfpatfylpramehrvgeesvrfdevcvvadrksfsllvecikgngyhftahpdwnvplrpdyldardevakklgltessklnglgtaas* |
| F | pB673 | meqkliseedleridghrqvvsnataqpgpllrpadmqaralqdlfdaqgvgvpvehalrmqavarqtntvfgirpverivttlieegfptkgfsvkgkssnwgpqagficvdqhlskredrdtaeirklnlavakgmdggaytqtdlrisrqrlaelvrnfglvadgvgpvrlltaqgpsgkryefearqepdglyrisrlgrseavqvlaspacglamtadydlflvapsieahgsggldarrntavrytplgakdplsedgfygredmargnitprtrqlvdalndclgrgehremfhhsddagnpgshmgdnfpatfylpramehrvgeesvrfdevcvvadrksfsllvecikgngyhftahpdwnvplrplfttkrklshdyldardevakklgltessklnglgtaas* |
| G | pB679 | meqkliseedleridghrqvvsnataqpgpllrpadmqaralqdlfdaqgvgvpvehalrmqavarqtntvfgirpverivttlieegfptkgfsvkgkssnwgpqagficvdqhlskredrdtaeirklnlavakgmdggaytqtdlrisrqrlaelvrnfglvadgvgpvrlltaqgpsgkryefearqepdglyrisrlgrseavqvlaspacglamtadydlflvapsieahgsggldarrntavrytplgakdplsedgfygredmargnitprtrqlvdalndclgrgehremfhhsddagnpgshmgdnfpatfylpramehrvgeesvrfdevcvvadrksfsnfqqvainasyraslndkwnsgldsplfttkrklshdyldardevakklgltessklnglgtaas* |
| H | pB680 | meqkliseedleridghrqvvsnataqpgpllrpadmqaralqdlfdaqgvgvpvehalrmqavarqtntvfgirpverivttlieegfptkgfsvkgkssnwgpqagficvdqhlskredrdtaeirklnlavakgmdggaytqtdlrisrqrlaelvrnfglvadgvgpvrlltaqgpsgkryefearqepdglyrisrlgrseavqvlaspacglamtadydlflvapsieahgsggldarrntavrytplgakdplsedgfygredmargnitprtrqlvdalnd |

| | | |
|---|---|---|
| | | clgrgehremfhhsddagnpgshmgdnfpatfylprhffdddglgegkgsiqtyfnvneqgavviqnpqefsnfqqvainasyraslndkwnsgldsplfttkrklshdyldardevakklgltessklnglgtaas* |
| I | pB682 | meqkliseedleridghrqvvsnataqpgpllrpadmqaralqdlfdaqgvgvpvehalrmqavarqtntvfgirpverivttlieegfptkgfsvkgkssnwgpqagficvdqhlskredrdtaeirklnlavakgmdggaytqtdlrisrqrlaelvrnfglvadgvgpvrlltaqgpsgkryefearqepdglyrisrlgrseavqvlaspacglamtadydlflvapsieahgsggldarrqpltweqwkesvtyedlspkykarydnqalyekqdgaslgnitprtrqlvdalndclgrgehremfhhsddagnpgshmgdnfpatfylpramehrvgeesvrfdevcvvadrksfsllvecikgngyhftahpdwnvplrplfttkrklshdyldardevakklgltessklnglgtaas* |
| J | pB683 | meqkliseedleridghrqvvsnataqpgpllrpadmqaralqdlfdaqgvgvpvehalrmqavarqtntvfgirpverivttlieegfptkgfsvkgkssnwgpqagficvdqhlskredrdtaeirklnlavakgmdggaytqtdlrisrqrlaelvrnfglvadgvgpvrlltaqgpsgkryefearqepdglyrisrlgrseavqvlaspacglamtadydlflvapsieahgsggldarrqpltweqwkesvtyedlspkykarydnqalyekqdgaslgnitprtrqlvdalndclgrgehremfhhsddagnpgshmgdnfpatfylpramehrvgeesvrfdevcvvadrksfsnfqqvainasyraslndkwnsgldsplfttkrklshdyldardevakklgltessklnglgtaas* |
| K | pB684 | meqkliseedleridghrqvvsnataqpgpllrpadmqaralqdlfdaqgvgvpvehalrmqavarqtntvfgirpverivttlieegfptkgfsvkgkssnwgpqagficvdqhlskredrdtaeirklnlavakgmdggaytqtdlrisrqrlaelvrnfglvadgvgpvrlltaqgpsgkryefearqepdglyrisrlgrseavqvlaspacglamtadydlflvapsieahgsggldarrqpltweqwkesvtyedlspkykarydnqalyekqdgaslgnitprtrqlvdalndclgrgehremfhhsddagnpgshmgdnfpatfylprhffdddglgegkgsiqtyfnvneqgavviqnpqefsnfqqvainasyraslndkwnsgldsplfttkrklshdyldardevakklgltessklnglgtaas* |
| L | pB717 | meqkliseedleridghrqvvsnataqpgpllrpadmqaralqdlfdaqgvgvpvehalrmqavarqtntvfgirpverivttlieegfptkgfsvkgkssnwgpqagficvdqhlskredrdtaeirklnlavakgmdggaytqtdlrisrqrlaelvrnfglvadgvgpvrlltaqgpsgkryefearqepdglyrisrlgrseavqvlaspacglamtadydlflvaptygdlgpqdkvkqpltweqwkesvtyedlspkykarydnqalyekqdgaslgmvsdrlkrtrqlvdalndclgrgehremfhhsddagnpgshmgdnfpatfylpramehrvgeesvrfdevcvvadrksfsllvecikgngyhftahpdwnvplrpsfqealdffqrkvnpgtaas* |
| M | pB719 | meqkliseedleridghrqvvsnataqpgpllrpadmqaralqdlfdaqgvgvpvehalrmqavarqtntvfgirpverivttlieegfptkgfsvkgkssnwgpqagficvdqhlskredrdtaeirklnlavakgmdggaytqtdlrisrqrlaelvrnfglvadgvgpvrlltaqgpsgkryefearqepdglyrisrlgrseavqvlaspacglamtadydlflvaptygdlgpqdkvkqpltweqwkesvtyedlspkykarydnqalyekqdgaslgmvsdrlkrtrqlvdalndclgrgehremfhhsddagnpgshmgdnfpatfylpramehrvgeesvrfdevcvvadrksfsllvecikgngyhftahpdwnvplrpdyldardevakklgltessklnglgtaas* |
| N | pB729 | meqkliseedleridghrqvvsnataqpgpllrpadmqaralqdlfdaqgvgvpvehalrmqavarqtntvfgirpverivttlieegfptkgfsvkgkssnwgpqagficvdqhlskredrdtaeirklnlavakgmdggaytqtdlrisrqrlaelvrnfglvadgvgpvrlltaqgpsgkryefearqepdglyrisrlgrseavqvlaspacglamtadydlflvaptygdlgpqdkvkqpltweqwkesvtyedlspkykarydnqalyekqdgaslgmvsdrlkelkdvintslgrtdglemvhhgaddanpyavmadnfpatffvpkhffdddglgegkgsiqtyfnvneqgavviqnpqefsnfqqvainasyraslndkwnsgldsplfttkrklshdyldardevakklgltessklnglgtaas* |
| O | pB730 | meqkliseedlegynygqalqeaqldiatmkprqrvtanelqlgddnaitnavtseqeatpnqdgshktyqsrdlvlepiqhpksielgmpevdqsvlaevaerenviigvrpvdeksksliaskmysskglfvkakssdwgpmsgfipvdqsfakasarrdlekfneyaeqsilsgnavsanlylnqvrieelvskyesltpleldvdsgmykttatngdqtipfflnkvtvddkelwqvhylregelapfkvigdpvskqpmtadydlltvmysieahgsggldarrntavrytplgakdplsedgfygredmargnitprtrqlvdalndclgrgehremfhhsddagnpgshmgdnfpatfylpramehrvgeesvrfdevcvvadrksfsllvecikgngyhftahpdwnvplrpsfqealdffqrkvnp* |

### Table S4. Overview of the protocol used for the molecular dynamics simulations.

| Step | Time | Positional restraints | Restraints force constant | Temperature control | Pressure control | Additional restraints |
|---|---|---|---|---|---|---|
| **Free ExoY** | | | | | | |
| Heating | 0.072 ns | All heavy atom excluding water and ions | 5 kcal/mol/A$^2$ | Langevin, t=50 - 300 K Damping = 1 ps$^{-1}$ | None | None |
| Equilibration 1 | 0.928 ns | All heavy atom excluding water and ions | 5 kcal/mol/A$^2$ | Langevin, t=300 K Damping = 1 ps$^{-1}$ | Nosé-Hoover Langevin, p= 1 atm Period = 100 fs Decay = 50 fs | None |
| Equilibration 2 | 2.5 ns | All heavy atom excluding water and ions | 2 kcal/mol/A$^2$ | Langevin, t=300 K Damping = 1 ps$^{-1}$ | Nosé-Hoover Langevin, p= 1 atm Period = 50 fs Decay = 25 fs | None |
| Equilibration 3 | 2.5 ns | Protein backbone | 2 kcal/mol/A$^2$ | Langevin, t=300 K Damping = 1 ps$^{-1}$ | Nosé-Hoover Langevin, p= 1 atm Period = 50 fs Decay = 25 fs | None |
| Equilibration 4 | 5 ns | Protein backbone except aa. 107 – 116, 317 - 326 | 2 kcal/mol/A$^2$ | Langevin; 300 K Damping = 1 ps -1 | Nosé-Hoover Langevin, p= 1 atm Period = 50 fs Decay = 25 fs | None |
| Equilibration 5 | 5 ns | None | - | Langevin, t=300 K Damping = 1 ps$^{-1}$ | Nosé-Hoover Langevin, p= 1 atm Period = 50 fs Decay = 25 fs | None |
| Production | 400 ns | None | - | Langevin, t=300 K Damping = 1 ps$^{-1}$ | Nosé-Hoover Langevin, p= 1 atm Period = 50 fs Decay = 25 fs | None |
| **F-actin ExoY** | | | | | | |
| Heating | 0.072 ns | All heavy atom excluding water and ions | 5 kcal/mol/A$^2$ | Langevin, t=50 - 300 K Damping = 1 ps$^{-1}$ | None | Mg coordination d = 2Å K = 50 kcal/mol/A$^2$ |
| Equilibration 1 | 0.928 ns | All heavy atom excluding water and ions | 5 kcal/mol/A$^2$ | Langevin, t=300 K Damping = 1 ps$^{-1}$ | Nosé-Hoover Langevin, p= 1 atm Period = 100 fs Decay = 50 fs | Mg coordination d = 2Å K = 50 kcal/mol/A$^2$ |
| Equilibration 2 | 2.5 ns | All heavy atom excluding water and ions | 2 kcal/mol/A$^2$ | Langevin, t=300 K Damping = 1 ps$^{-1}$ | Nosé-Hoover Langevin, p= 1 atm Period = 50 fs Decay = 25 fs | Mg coordination d = 2Å K = 50 kcal/mol/A$^2$ |
| Equilibration 3 | 2.5 ns | Protein backbone | 2 kcal/mol/A$^2$ | Langevin, t=300 K Damping = 1 ps$^{-1}$ | Nosé-Hoover Langevin, p= 1 atm Period = 50 fs Decay = 25 fs | Mg coordination d = 2Å K = 50 kcal/mol/A$^2$ |
| Equilibration 4 | 5 ns | Protein backbone except: actin aa. 1 – 4; ExoY aa. 107 – 116, aa. 317 - 326 | 2 kcal / mol A2 | Langevin, t=300 K Damping = 1 ps$^{-1}$ | Nosé-Hoover Langevin, p= 1 atm Period = 50 fs Decay = 25 fs | Mg coordination d = 2Å K = 50 kcal/mol/A$^2$ |
| Equilibration 5 | 5 ns | None | - | Langevin, t=300 K Damping = 1 ps$^{-1}$ | Nosé-Hoover Langevin, p= 1 atm Period = 50 fs Decay = 25 fs | Mg coordination d = 2Å K = 50 kcal/mol/A$^2$; Orientation restraint (orientation colvar) Restraint on every 5th CA of the last two protomers of each of actin's end K = 1000 |
| Production | 400 ns | None | - | Langevin, t=300 K Damping = 1 ps$^{-1}$ | Nosé-Hoover Langevin, p= 1 atm Period = 50 fs Decay = 25 fs | Mg coordination d = 2Å K = 50 kcal/mol/A$^2$ Orientation restraint (orientation colvar) Restraint on every 5th CA of the last two protomers of each of actin's end K = 400 |

**Movie S1. Molecular dynamics simulations of free PaExoY and in complex with F-actin.** For guidance, the starting structure of PaExoY is shown as translucent. For the F-actin complexes, we show only the initial structure of the filament as surface.

**Movie S2. Mechanism of activation of PaExoY upon binding to F-actin.**
